# Monitoring monomer-specific acyl-tRNA levels in cells with PARTI

**DOI:** 10.1101/2025.01.14.633079

**Authors:** Meghan A. Pressimone, Carly K. Schissel, Isabella H. Goss, Cameron V. Swenson, Alanna Schepartz

## Abstract

We describe a new assay that reports directly on the acylation state of a user-chosen tRNA in cells. We call this assay 3-Prime Adenosine-Retaining Aminoacyl-tRNA Isolation (PARTI). It relies on high-resolution mass spectrometry identification of the acyl-adenosine species released upon RNase A cleavage of isolated cellular tRNA. Here we develop the PARTI workflow and apply it to understand three recent observations related to the cellular incorporation of non-α-amino acid monomers into protein: (1) the origins of the apparent selectivity of translation with respect to β^2^-hydroxy acid enantiomers; (2) the activity of PylRS variants for benzyl derivatives of malonic acid; and (3) the apparent inability of *N*-Me amino acids to function as ribosome substrates in living cells. Using the PARTI assay, we also provide direct evidence for the cellular production of 2’,3’-diacylated tRNA in certain cases. The ease and simplicity of the PARTI workflow should benefit ongoing efforts to study and improve the cellular incorporation of non-α-amino acid monomers into proteins.

**GRAPHICAL ABSTRACT:** 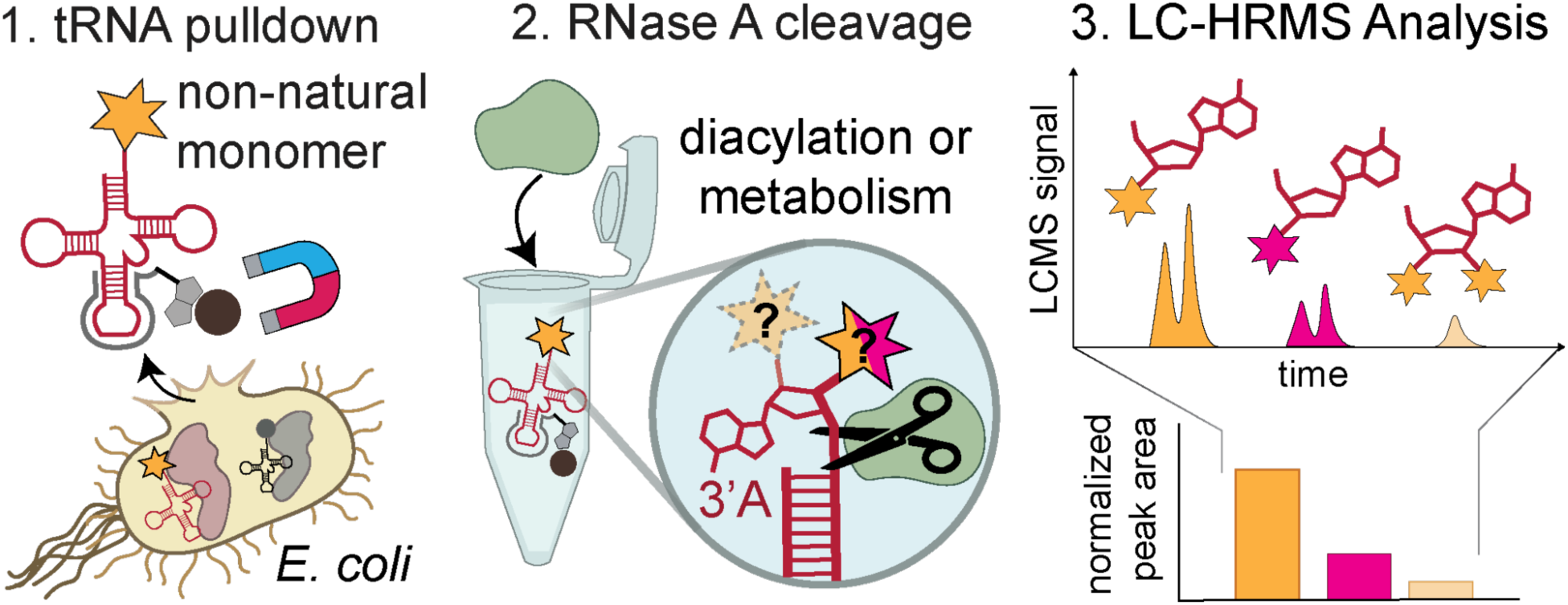

## INTRODUCTION

There is widespread interest in the cellular biosynthesis of genetically encoded materials containing one or more non-α-amino acid monomers (1). Even monomers that differ from α-amino acids by a single atom, such as α-hydroxy acids, can generate proteins with unique properties. Examples include discrete amide to ester substitutions that probe ion channel function (2) or the contributions of backbone H-bonds to protein stability (3), and others that support intramolecular rearrangements to generate expanded or altered backbones (4, 5). Over the past few years, a small number of non-α-amino acids, notably β^2^-hydroxy and β^3^-amino acids, have been introduced into proteins in cells using native machinery or orthogonal aminoacyl tRNA synthetase (aaRS)/tRNA pairs and stop codon suppression (6–8). Yet even with highly active or evolved aaRS enzymes, protein yields are low, unpredictable, or both (7, 8). Although ribosomal translation is complex and multi-step, and myriad events could contribute to low or unpredictable protein yields (3, 9–11), the level of tRNA aminoacylation is essential. A robust assay that reports directly on tRNA acylation levels in cells would streamline efforts to optimize the cellular incorporation of non-α-amino acid monomers into proteins. An assay that reports simultaneously on monomer identity would provide an even more accurate snapshot of the state of tRNA acylation and thus the specificity of the corresponding aaRS.

The challenge in the development of an assay that reports simultaneously on cellular tRNA acylation and monomer identity is that expression of an aaRS/tRNA pair in the presence of an aaRS substrate can result in multiple different acyl-tRNA products (**Figure 1A**). These products include the expected 3’-monoacylated tRNA as well as those carrying a second acyl group on the 2’-hydroxyl groups as observed *in vitro* (7, 12, 13), and others that are misacylated with an incorrect or metabolized version of the aaRS substrate as observed in cells using protein translation as a proxy (3, 9, 14). Few methods directly capture the *in vivo* acylation level of a single tRNA of interest. Of those that do, most rely on differentiating acylated and unreacted tRNA populations using periodate oxidation and fail to distinguish between different potential acyl-tRNA products (15–18).

**Figure 1:**
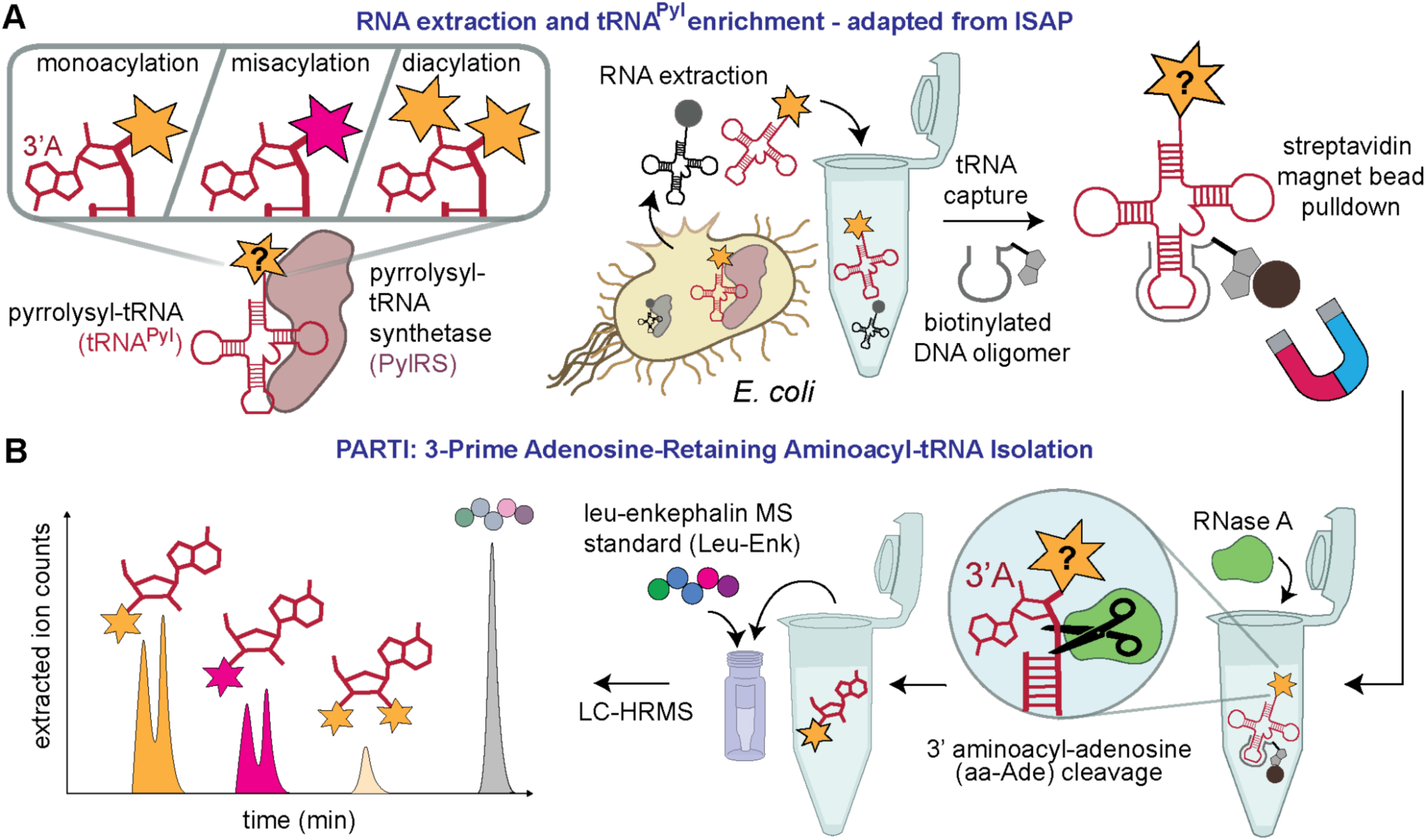
3-Prime Adenosine-Retaining aminoacyl-tRNA Isolation (PARTI) reports on monomer-specific tRNA acylation in cells. **(A)** Schematic showing three states of tRNA – monoacylated, misacylated, or diacylated – that can exist in *E. coli* expressing an aaRS/tRNA pair and supplemented with substrate. As per ISAP, the tRNA to be analyzed is selectively captured from total cellular RNA using a biotinylated complementary DNA oligonucleotide and sequestered using streptavidin-coated magnetic beads. **(B)** In the PARTI workflow, the sequestered tRNA population is treated with RNase A to release the 3’ aminoacyl adenosine (aa-Ade) which is then detected using LC-HRMS and quantified using a peptide standard, leucine-enkephalin (Leu-Enk).

One recently reported method that provides a partial snapshot of the cellular tRNA acylation state, including the identity of the acylated monomer, is referred to as isoacceptor-specific affinity purification (ISAP) (19). Here, total RNA is first isolated from cells grown in the presence of an aaRS substrate and the tRNA of interest is extracted from total cellular RNA using a complementary biotinylated DNA oligonucleotide (**Figure 1B**). The population of acyl-tRNAs is subsequently hydrolyzed and the monomer detected by mass spectrometry (19).

Although ISAP is highly sensitive, the post-capture hydrolysis step demands extensive washing to remove contaminating amino acids and leverages radiolabeling to detect low levels of misacylation, which is challenging for non-canonical monomers not available in radiolabeled form. Additionally, because it relies on hydrolysis of the acylated tRNA to release the subsequently detected monomer, it conflates tRNA populations that are mono- and diacylated. As a result, ISAP overestimates the acylation yield, and hence the activity, of any monomer that doubly acylates tRNA in cells, as has been observed for multiple backbone-modified monomers *in vitro* (7, 9, 12). Complicating matters further is the fact that tRNAs acylated with some non-α-amino acids resist hydrolysis (20), which underestimates acylation yield and activity.

Here we describe a variation of ISAP that overcomes these limitations to provide a detailed snapshot of cellular tRNA acylation. Rather than relying on the detection of monomers after acyl-tRNA hydrolysis, here we detect the acylated 3’-adenosine using high resolution mass spectrometry (HRMS) after enzymatic cleavage of the terminal 3’-adenosine using RNase A (**Figure 1B**) (9, 21). Detecting the 3’-adenosine by mass spectrometry unambiguously identifies the acylated species and whether the tRNA has been diacylated on both the 2’ and 3’ position.

We refer to this new assay as PARTI, for 3-<u>P</u>rime <u>A</u>denosine-<u>R</u>etaining Aminoacyl-tRNA <u>I</u>solation (**Figure 1B**). Here we develop a robust PARTI workflow and apply it to address three outstanding questions related to the cellular synthesis of proteins containing one or more non-α-amino acid monomers: the enantioselectivity of pyrrolysyl tRNA synthetase (PylRS) in cells with respect to β^2^-hydroxy acid substrates (7), the cellular activity of PylRS variants with respect to benzyl malonate substrates (9), and the metabolic fate of *N*-methyl (*N-*Me) amino acids in cells, which have been introduced into peptides and proteins, but only in *in vitro* translation mixtures and cell extracts (22–30). We anticipate that PARTI will aid the analysis of cellular tRNA acylation status and benefit ongoing efforts to expand the proteome to include diverse backbone-modified monomers.

## METHODS

### GENERAL METHODS

Note: Procedures involving RNA were carried out using RNase-free materials and technique, which includes decontaminating the work area with RNase FREE (Apex Bio) or a similar product.

### Transcription and purification of *in vitro* tRNAs

DNA templates used to transcribe either *E. coli* tRNA^Phe^ (*Ec*tRNA^Phe^) or *Methanomethylophilus alvus* tRNA^Pyl^ (*Ma*tRNA^Pyl^) were generated by annealing and extending the single-stranded DNA oligonucleotide pairs PheT-Fwd and PheT-Rev or PylT-Fwd and PylT-Rev (2 mM; **Supplementary Information, Table S1**) using OneTaq 2x Master Mix (NEB) according to the manufacturer protocol.

Both *Ec*tRNA^Phe^ and *Ma*tRNA^Pyl^ were transcribed *in vitro* using the TranscriptAid Kit (NEB) according to the manufacturer protocol. Transcription reactions of a final volume of 200 µL were divided into eight 25 µL aliquots and incubated at 37 °C for 4 h. Then, the eight 25 µL reactions were pooled, 20 µL DNase I (NEB) was added, and the reaction was incubated at 37 °C for 1 h followed by purification using gel electrophoresis and ethanol precipitation as described (31).

The tRNA was then analyzed for purity using intact tRNA LC-MS as described (32). A representative analysis of *Ec*tRNA^Phe^ is shown in Supplementary Information, Figure S1.

### Cell Transformations

Chemically competent cells in 100 µL aliquots were transformed by incubating with 100 ng plasmid DNA on ice for 30 min followed by a 90 sec heat-shock at 42 °C. The cells were allowed to recover on ice for 2 min, supplemented with 800 µL LB, grown at 37 °C for one hour with shaking, and applied to a LB/agar plate prepared with the appropriate antibiotic.

### Purification of aminoacyl-tRNA synthetases

*M. alvus* PylRS (*Ma*PylRS) was purified from BL21-Gold (DE3)pLysS chemically competent cells and analyzed as described (9) except that TALON resin (Takara Bio) was used rather than Ni-NTA agarose resin. The yield was 170 mg/L

*Ec*PheRS was purified from BL21-Gold (DE3)pLysS chemically competent cells using the same protocol as for *Ma*PylRS (9) with the following changes. The plasmid used to express *Ec*PheRS was derived from pET-32a and contained the sequences of the α and β subunits of *Ec*PheRS; a His_6_-SUMO tag was encoded at the N-terminus of the αsubunit. Following purification with TALON resin, the protein concentration was determined using a NanoDrop ND-1000 device (Thermo Scientific). The His_6_-SUMO tag was removed by adding the purified protein and SUMO protease purified previously from BL21-Gold (DE3) cells transformed with a pET-32a plasmid containing the sequence for SUMO protease with a His_6_ tag encoded at the N-terminus (33) in an 8:1 molar ratio to Snakeskin™ dialysis tubing (10K MWCO, Thermo) equilibrated with Storage Buffer (100 mM HEPES-K (pH 7.2), 100 mM NaCl, 10 mM MgCl2, 4 mM dithiothreitol (DTT), 20% (v/v) glycerol) and incubated for 16 h at 4 °C. The protein was then concentrated using an Amicon® Ultra Centrifugal Filter, 10 kDa MWCO and the buffer exchanged to 50 mM sodium phosphate (pH 7.4), 500 mM NaCl, 20 mM β-mercaptoethanol using a PD-10 desalting column packed with Sephadex G-25 resin (Cytiva). *Ec*PheRS was then isolated from the His_6_-SUMO tag by incubating the sample with TALON resin for 45 min at 4°C and collecting the flowthrough containing *Ec*PheRS. *Ec*PheRS was analyzed by LC-HRMS as described (9) (**Supplementary Information, Figure S1**). Protein concentration was determined by Pierce™ Bradford Assay (Thermo Scientific). The yield was 88 mg/L.

### *In vitro* tRNA acylation and analysis

Acylation of *in vitro*-transcribed and purified *Ma*tRNA^Pyl^ and *Ec*tRNA^Phe^ was performed as previously described (9) using 25 µM tRNA and 1.5-12.5 µM of the requisite aaRS for 2-4 h, as specified per each experiment. Purified *Ma*PylRS was used to acylate *Ma*tRNA^Pyl^ with 10 mM of a given substrate: L-α-amino-Boc-lysine, (BocK) (Sigma Aldrich), L-α-methyl-amino-Boc-lysine (*N*-Me-BocK), (S)-6-((tert-Butoxycarbonyl)amino)-2-hydroxyhexanoic acid (OH-BocK) (Accela Chembio), (R)-6-((tert-butoxycarbonyl)amino)-2-(hydroxymethyl)hexanoic acid (*(R)-*β_2_-OH-BocK) (Enamine), or *(S)*-6-((tert-butoxycarbonyl)amino)-2-(hydroxymethyl)hexanoic acid (*(S)-*β_2_-OH-BocK) (Enamine). Purified *Ec*PheRS was used to acylate *Ec*tRNA^Phe^ with 10 mM L-phenylalanine (Phe) (Frontier Scientific). Samples were analyzed by intact tRNA LC-MS and RNase A/LC-HRMS as described (9, 32). The major ion (m/z), corresponding peak area, and percent acylation for each replicate can be found in **Supplementary Information, Table S3**.

### Statistical analysis

Either a one-way ANOVA or two-tailed t test was performed for statistical comparisons. Exact parameters and results can be found in **Supplementary Information, Table S2**.

## 2. PARTI Workflow

### Design of biotinylated DNA capture oligonucleotide o-Pyl

The sequence of the capture oligonucleotide for tRNA^Pyl^, o-Pyl, was designed to match the length and predicted T_m_ of the established capture oligonucleotide for tRNA^Phe^ (19), referred to here as o-Phe, with slight changes related to potential differences in modified bases. It is not known whether *Ma*tRNA^Pyl^ contains modified bases, but the homolog from *M. barkeri* is proposed to have a 4-thiouridine at position 8 and 1-methyl pseudouridine at position 50 (34). To avoid these positions, the o-Pyl sequence used for PARTI was complementary to residues 27 to 49 and encompasses the full anticodon loop and half the anticodon and T stems (**Supplementary Information, Table S1**), with a total of 5 fewer bases (26 versus 31 bases) than o-Phe and a T_m_ predicted to be 2 °C lower (62.8 °C versus 64.5 °C), as predicted using Benchling software.

While the capture oligonucleotide used in ISAP was biotinylated on the 3’ end, o-Phe and o-Pyl as used in this work are biotinylated on the 5’ end and were purchased from IDT with this modification.

**Cell Growth**. The lac-inducible plasmids pMEGA-PylRS-PylT or pMEGA-FRSA-PylT (9) were transformed into either DH5α, Top10 or C321.ΔA.exp (C321) (35) chemically competent *E. coli* cells. The cells were then plated on a LB/agar plate prepared with 0.1 mg/mL spectinomycin and incubated overnight at 37 °C. A single colony was chosen and used to inoculate 10 mL of LB containing spectinomycin (0.1 mg/mL) and the cells grown with shaking at 200 rpm for 16 h. This overnight culture was used to inoculate subsequent growths as described below.

*For isolation of tRNA^Phe^:* 500 mL of LB containing 0.1 mg/mL spectinomycin was inoculated with 5 mL of the overnight culture and grown at 37 °C with shaking at 200 rpm until the OD_600_ reached 0.4–0.6. Expression of tRNA^Pyl^ and PylRS was *not* induced. Cultures were pelleted by centrifugation using a Beckman Coulter Allegra®X-14R Benchtop Centrifuge using a Beckman SX4750 Swinging Bucket Rotor, frozen at -80 °C and stored for up to a month before RNA extraction as described below.

*For isolation of tRNA^Pyl^:* 50 mL (unless otherwise noted) of LB containing 0.1 mg/mL spectinomycin was inoculated with 5 mL of the overnight culture, supplemented with aaRS substrate (final concentration 0.1-1 mM), and grown at 37 °C with shaking at 200 rpm. Cultures were grown until the OD_600_ reached 0.4–0.6 then were supplemented with IPTG to a final concentration of 1 mM to induce the expression of aaRS and tRNA^Pyl^ genes. The induced cultures were grown for 3 h at 37 °C with shaking at 200 rpm, then pelleted by centrifugation using a Beckman Coulter Allegra®X-14R Benchtop Centrifuge using a Beckman SX4750 Swinging Bucket Rotor and frozen at -80 °C. Pellets were stored for up to a month before RNA extraction as described below.

### Total RNA Extraction

Samples were maintained on ice during all extraction steps. TRIzol™ Reagent (Thermo Scientific) was used and the extraction executed according to the manufacturer’s protocol. 3 mL Trizol was added to pellets originating from 500 mL uninduced log phase cells grown for tRNA^Phe^ pulldown and 1 mL Trizol was added to pellets originating from 50 mL cultures expressing tRNA^Pyl^. Samples were mixed by vortexing and incubated on ice for 15 min to lyse the cells. Then 0.2 mL chloroform was added for every 1 mL Trizol, and a liquid-liquid extraction was performed. The aqueous layer containing total RNA was transferred to new 1.7 mL Eppendorf tubes and 0.5 mL isopropanol was added for every 1 mL Trizol. Samples were mixed by inverting the tubes then incubated on ice for 15 min to precipitate the RNA. Samples were centrifuged in a Eppendorf™ 5425 Centrifuge for 30 min at 21,000 relative centrifugal force (rcf) and 4 °C to pellet the RNA. The supernatant was discarded and the pellet was washed with 500 µL 70% ethanol and centrifuged for 10 min at 21,000 rcf and 4 °C. The supernatant was discarded and the RNA pellet dried at RT for 10 min. The pellet was either frozen at -80 °C or resuspended immediately. When ready to use, dried pellets were resuspended in 200 µL RNase free water (Milli Q). RNA concentration was determined by measuring the absorbance at 260 nm using a NanoDrop ND-1000 device (Thermo Scientific) and samples were kept on ice before tRNA isolation using o-Phe or o-Pyl.

### tRNA isolation using o-Phe or o-Pyl

The following steps are adapted from Mohler *et al.* (19), save for the RNA quantity used. In a new 1.7 mL Eppendorf tube, 250 µg resuspended total RNA was mixed with 5 µL 100 µM biotinylated DNA oligomer (IDT) (500 pmol, final concentration 1 µM), 250 µL 4x saline-sodium citrate (SSC) (Invitrogen) pH 4.8 (final concentration 2X), and MilliQ RNase free water up to a final volume of 500 µL. The solution was incubated for 60 min at 50 °C on a Thermolyne Dri-Bath to generate the tRNA-DNA hybrid and then cooled to RT. While the tRNA-DNA hybrid solution was cooling to RT, 0.5 mg Streptavidin MagneSphere® Paramagnetic Particles (SA-PMPs) (Promega) (500 µL of a 1 mg/mL resuspension) were added to a fresh 1.7 mL tube. The maximum capacity of 0.5 mg of streptavidin-coated magnetic beads is estimated to be 625 pmol according to the Product Technical Bulletin. The tube was secured in a magnetic rack (MagRack 6, Cytiva) and the supernatant was removed carefully to not include any streptavidin-coated magnetic beads. The beads were washed 3 times with 300 µL 2x SSC pH 4.8 and the supernatant discarded before the tRNA-DNA hybrid solution was added to the beads and rotated for 10 min. The supernatant was then discarded, and the beads were washed 6 times with 300 µL 2x SSC pH 4.8. Finally, the beads were resuspended in 10 µL 2x SSC pH 4.8.

### RNase A cleavage and addition of MS standard

Note: When using RNase A, special care must be taken to avoid contamination of surfaces, pipettes, etc. Surfaces should be decontaminated with RNase FREE (Apex Bio) or a similar product. 11 µL 1.51 U/µL RNase A (VWR) dissolved in 200 mM NaOAc was added to the resuspended beads prepared as described above, and the mixture was incubated for 10 min at RT (21). 2.2 µL 50% TCA was added and the sample was incubated for an additional 10 min to halt the RNase A cleavage reaction (21). The samples were centrifuged for 10 min at 21,000 rcf and RT and a 20 µL aliquot of the supernatant was transferred to a new 1.7 mL tube and supplemented with 2 µL of a 40 ng/µL solution of leucine-enkephalin (Leu-Enk) (Waters). This solution diluted 1:10 with Mobile Phase A (0.1% formic acid in water) into a SureSTART™ 0.3 mL Glass Screw Top Microvial (Thermo Scientific) before LC-HRMS analysis as described below. Because the streptavidin-coated magnetic beads can retain an inconsistent volume of 2x SSC from wash steps during tRNA isolation, the remaining volume of supernatant beyond the 20 µL used for LC-HRMS was recorded for quantitatively determining A_norm_. The equation for determining A_norm_ is provided below.

### LC-HRMS and Analysis

1 µL of the sample prepared as described above was analyzed by LC-HRMS as described (9), with the following modifications. Chromatography was performed at a flow rate of 0.5 mL/min using mixtures of Mobile Phase B (100% acetonitrile) and Mobile Phase A (0.1% formic acid in water). For each injection at t = 0, the eluent was initially held at 4% Mobile Phase B for 1.89 min. The amount of Mobile Phase B was then increased linearly from 4 to 40% over 3.11 min, then from 40 to 100% Mobile Phase B over 2 min, from 100 to 4% Mobile Phase B over 2 min, and finally held at 4% Mobile Phase B for 0.5 min to complete the chromatographic run. Mass spectrometry data was collected as described (9) between 1.4 and 7 min, with m/z ranging 100-3000 recorded.

A_norm_ values were determined as follows: First, the presence of free Ade or a given mono- or diacyl adenosine species was confirmed by generating an extracted ion chromatogram (EIC) of the expected exact [M+H] to five decimal places, determined using ChemDraw software, over a symmetric range of 100 ppm. Free Ade yielded one peak. aa-Ade appeared as two EIC peaks corresponding to the 2′- and 3′-acyl-adenosine products for chiral monomers or as more peaks for prochiral monomers. Di-aa-Ade, when observed, appeared as a single peak in the respective EIC. Chromatographic peaks containing the exact mass (error < 5 ppm) as well as the expected mass envelope were counted. Then, for quantification, new EICs were generated by extracting over the m/z range of the monoisotopic peak present in the sample. This process was repeated for Leu-Enk in the sample. The integrated areas of each analyte peak were recorded and the following equation was used to generate the corresponding A_norm_ value:

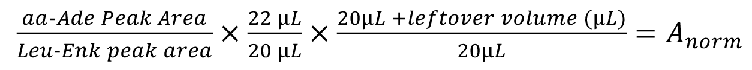

## 3. METHODS IN SUPPORT OF INDIVIDUAL FIGURES

### Methods in Support of Figure 2

*In vitro* acylation reactions were performed using 10 mM Phe, 25 µM tRNA^Phe^ and 1.5 µM purified *E. coli* PheRS in a final volume of 25 µL for 2.5 h at 37 °C and was otherwise the same as described in the methods for *in vitro* acylation. From a 15 µL aliquot of the reaction, Phe-tRNA^Phe^ was isolated by phenol chloroform extraction and ethanol precipitation then resuspended in water. Intact-tRNA LC-MS was carried out as described previously with 10 pmol of sample (32). The remaining 250 pmol of the tRNA^Phe^ acylation in 10 µL was treated with RNase A as described. LC-HRMS was carried out as described except for the length of run time and MS collection window. Chromatography was carried out with mobile phase B at 4% for 1.89 min followed by a linear gradient from 4-40% over 1.75 min. Then, mobile phase B underwent a gradient from 40-100% over 0.56 min and a subsequent gradient from 100-4% over 0.98 min.

Mobile phase B was held at 4% for an additional 1.12 min. Mass spectrometry data were collected for the entire duration of the LC-HRMS run.

*In vivo* Phe-tRNA^Phe^ was purified first by growing *E. coli* DH5α cells transformed with a pET32a plasmid containing a carbenicillin resistance gene and a T7-promoted *Ma*PylRS gene; the latter was not expressed because DH5α cells do not express T7 RNA polymerase (36). The PARTI protocol was carried out as described but with 95 µg starting total RNA. LC-HRMS analysis was performed in the same manner as for the i*n vitro* Phe-tRNA^Phe^ sample described in this section.

### Methods in Support of Figure 3

PARTI was carried out as described for purification of tRNA^Phe^ from *E. coli* DH5α cells transformed with, but not expressing, pMEGA-PylRS-PylT.

### Methods in Support of Figure 4

An *in vitro* acylation was carried out as described with 25 µM *in vitro*-transcribed and purified tRNA^Pyl^, 10 µM purified *M. alvus* PylRS, and 10 mM BocK in 50 µL for 2 hours at 37°C. BocK-tRNA^Pyl^ was then purified using the RNA Clean & Concentrate Kit (Zymo) and the concentration was determined using a NanoDrop ND-1000 device (Thermo Scientific). Then, PARTI was carried out using o-Pyl as described with 500 pmol total tRNA^Pyl^. In parallel, 10 pmol of total tRNA^Pyl^ was analyzed by intact RNA LC-MS to determine the acylation yield.

### Methods in Support of Figure 5

PARTI was carried out as described for purification of tRNA^Pyl^ from *E. coli* C321 cells transformed with pMEGAPylRS/PylT and incubated with no substrate or 0.1 mM either BocK, OH-BocK, (*S)*-β_2_-OH-BocK, or (*R)*-β_2_-OH-BocK.

### Methods in Support of Figure 6

PARTI was carried out as described for purification of tRNA^Pyl^ with the following modifications. 30 mL of LB containing 0.1 mg/mL spectinomycin was inoculated with 3 mL of an overnight culture of *E. coli* C321 cells transformed with pMEGA-FRSA/PylT and supplemented with either no substrate or 1 mM of 3-Bromo-L-phenylalanine (*m-*Br-Phe) (CombiBlocks), or of previously synthesized (9) 2-(3-trifluoromethyl)malonic acid (*m*-CF_3_-bma), (S)-2-amino-3-(3-bromophenyl)propanoic acid (*m*-Br-bma), or 2-(3-methylbenzyl)malonic acid.

### Methods in Support of Figure 7

PARTI was carried out as described for purification of tRNA^Pyl^ with *E. coli* Top10 cells transformed with pMega-PylRS/PylT were grown with either no substrate, 1 mM BocK, or 1 mM N-Me BocK.

### Expression, Purification, and LC-HRMS analysis of sfGFP-200TAG

The expression, purification, and LC-HRMS analysis of sfGFP-200TAG was performed as described (9) with the following changes. Chemically competent *E. coli* Top10 cells were transformed with sfGFP-200TAG and pMega-*Ma*PylRS. TB media was used for outgrowth during sfGFP-200TAG expression and cultures were supplemented with either 1 mM BocK or 1 mM *N*-Me BocK. Yields were 36 mg/L when expression was supplemented with 1 mM BocK and 8 mg/L when expression was supplemented with 1 mM *N*-Me BocK.

### Preparation of *N*-Me BocK

*N*-Me BocK was prepared from Fmoc-*N*-Me-Lys(Boc)-OH (Sigma Aldrich) using 20% piperidine in DCM as described (37). In brief, the reaction was stirred for 1.5 h and was determined to be complete by LC-MS. *N*-Me BocK was extracted three times with water and washed with DCM. The aqueous layer was flash frozen with dry ice/acetone and lyophilized for 5 days resulting in a white powder in quantitative yield.

### Small Molecule MS Analysis

Mixtures of *N*-Me BocK and BocK at 0.1 mg/mL (999:1, 99:1, and 90:10 I-Me BocK:BocK) were prepared and analyzed using a Waters SQD2 mass spectrometer fitted with a reverse-phase C18 column, a 200-800 nm UV/vis detector, a 300-1000 nm fluorescence detector. 1 µL of each mixture was injected. Chromatography was performed at a flow rate of 0.5 mL/min using mixtures of Mobile Phase B (0.1% formic acid in acetonitrile) and Mobile Phase A (0.1% formic acid in water). For each injection at t = 0, the eluent was initially set to 10% Mobile Phase B and was then increased linearly from 10% to 95% over 3.5 min, then decreased from 95% to 10% Mobile Phase B for 0.1 min, and finally held at 10% Mobile Phase B for 1.5 min to complete the chromatographic run. Mass spectrometry data with M/z ranging from 150-800 Da was collected in positive mode and in centroid form with the following parameters: cone voltage = 30 V, probe temperature = 20 °C, scan time = 0.25 s.

### Plate reader analysis of sfGFP expression

The protocol was carried out as described (9) with the following changes. Following transformation into *E. coli* Top10 or C321 cells and growth of starter cultures as described, 500 µL each culture was used to inoculate 25 mL TB supplemented with 0.1 mg/mL carbenicillin and 0.1 mg/mL spectinomycin in 250 mL baffled Erlenmeyer flasks. Cultures were grown to OD 0.6-0.8 before induction with IPTG (final concentration 1 mM). 180 µL induced culture was added to each well with 20 µL LB for no substrate control, or 20 µL substrate for final concentrations of either 1 mM *N*-Me BocK, 5 mM *N*-Me BocK, or 0-1 mM BocK. Substrates were dissolved in water and NaOH in equal molar amount to the monomer itself to ensure solubilization.

## RESULTS

### RNase A cleavage and LC-HRMS provide an accurate measure of tRNA acylation

To develop a robust PARTI workflow, we first set out to confirm that RNase A treatment of *in vitro*-acylated tRNA followed by LC-HRMS analysis provides an accurate measure of tRNA acylation. We began with a well-characterized reaction, that of *E. coli* PheRS with *E. coli* tRNA^Phe^ and phenylalanine, to generate Phe-tRNA^Phe^. Purified, *in vitro*-transcribed tRNA^Phe^ was acylated using purified PheRS and the products analyzed using both intact tRNA LC-MS (32) (**Figure 2A**) and RNase A digestion followed by LC-HRMS (21) (**Figure 2B**). Intact tRNA LC-MS of the acylation reaction mixture revealed two RNA products; one whose deconvoluted mass corresponds to unreacted tRNA^Phe^ and the other to tRNA^Phe^ monoacylated with Phe (Phe-tRNA^Phe^). Integration of the two peaks indicated an acylation yield of 61% (**Figure 2A**) (32).

**Figure 2:**
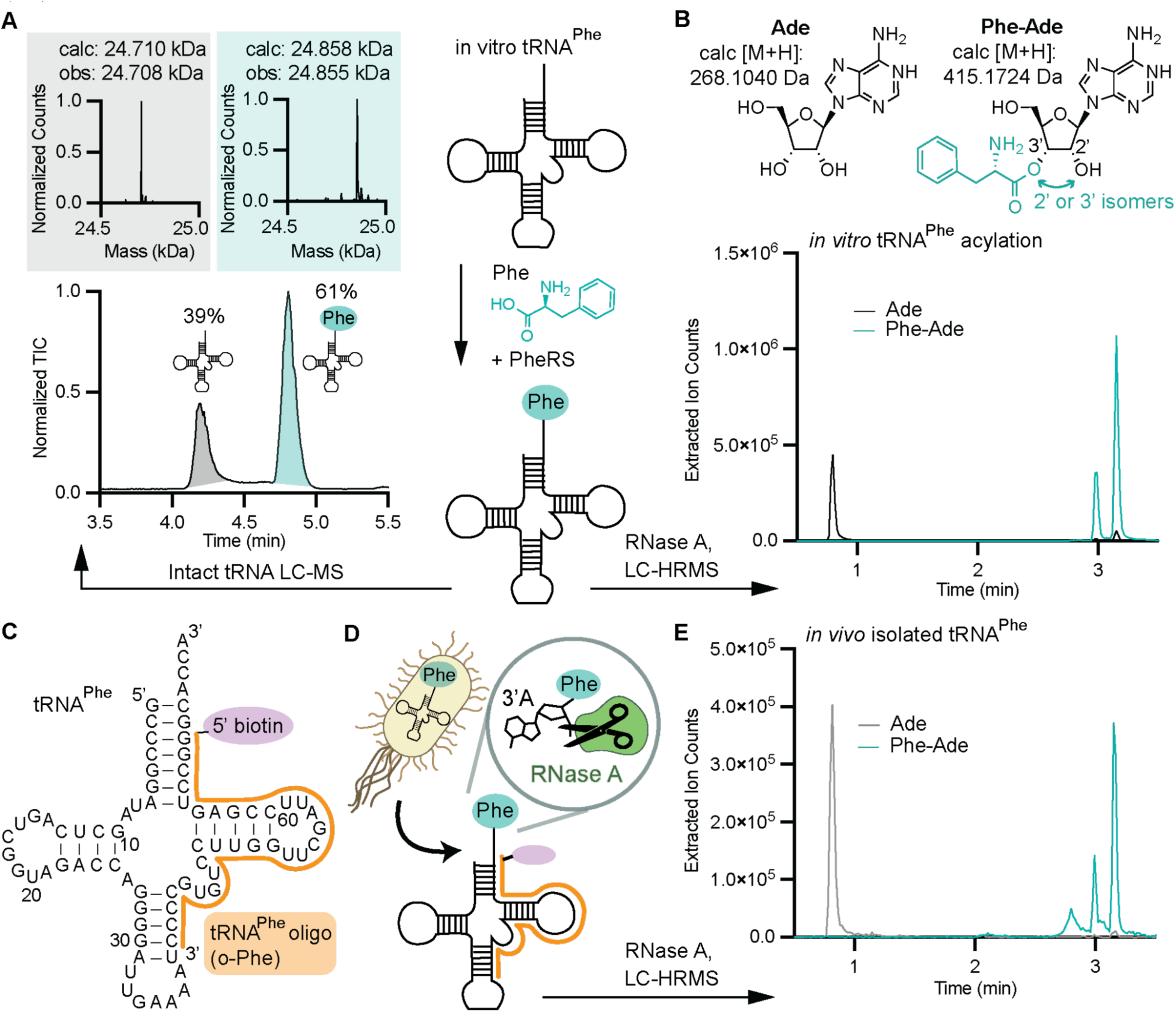
Validating the PARTI workflow *in vitro* and in cells using *E. coli* PheRS and tRNA^Phe^. **(A)** Intact tRNA LC-MS analysis of the products resulting from *in vitro* acylation of tRNA^Phe^ with Phe and PheRS. The total ion chromatogram (TIC) of the reaction mixture is shown along with the deconvoluted mass spectrum of each peak. The gray peak in the TIC corresponds to the deconvoluted spectrum in gray, with the expected and observed mass of unreacted tRNA^Phe^ shown. The teal peak in the TIC corresponds to the deconvoluted spectrum in teal, with the expected and observed mass of Phe-tRNA^Phe^. The yield of Phe-tRNA^Phe^ was determined by the ratio of the integrated areas of the major ion for Phe-tRNA^Phe^ and unreacted tRNA^Phe^ as previously described (32). **(B)** Extracted ion chromatograms (EICs) from LC-HRMS analysis of the products resulting from *in vitro* acylation of tRNA^Phe^ with Phe and PheRS following RNase A treatment. The trace in black is the EIC corresponding to the M+H for adenosine (Ade), whereas the trace in teal corresponds to the M+H for Phe-Ade, whose structure as a mixture of 2’ and 3’ isomers is shown. No diacyl-tRNA was detected. The EIC peak area for Ade and the summed areas of the two peaks due to Phe-Ade were used to calculate an acylation yield of 74%. **(C)** The secondary structure of *E. coli* tRNA^Phe^ bound to o-Phe, the biotinylated DNA capture oligonucleotide used to extract total tRNA^Phe^ from isolated total RNA. **(D)** Schematic of purification of tRNA^Phe^ from cells using o-Phe followed by RNase A cleavage and LC-HRMS analysis of the reaction products. **(E)** Overlaid traces showing the extracted ions for adenosine (Ade) in gray and phenylalanine-adenosine (Phe-Ade) in teal generated by applying the PARTI workflow to tRNA^Phe^ isolated from *E. coli.* The acylation yield is calculated to be 56% using the strategy described in **(B)**.

RNase A digestion of the acylation reaction followed by LC-HRMS also revealed two products: one whose mass corresponds to adenosine (Ade), the product expected from digestion of unreacted tRNA^Phe^, and the other (Phe-Ade) whose mass corresponds to the product expected from digestion of Phe-tRNA^Phe^ (**Figure 2B**). The ion chromatogram extracted for the mass of Phe-Ade (EIC) contains two peaks at ∼3 min which we assign to the anticipated mixture of 2’ and 3’-acylated products (9). The ion chromatogram extracted for the mass of Ade contains a single peak near 1 min and two small peaks at the same retention time as Phe-Ade which result from fragmentation of Phe-Ade during MS. Integrating the Phe-Ade and Ade EIC peaks indicates an acylation yield of 74%, which agrees reasonably well with the 61% yield determined by intact tRNA LC-MS. No diacyl-tRNA^Phe^ was observed under these conditions.

Next, we evaluated the extent of tRNA^Phe^ acylation in *E. coli* by sequestering the complete cellular population of acylated and unreacted cellular tRNA^Phe^ using the validated biotin-tagged capture oligonucleotide o-Phe (**Figure 2C** and **Supplementary Information, Table S1**) (19). Although *E. coli* contains two genes encoding tRNA^Phe^, the resulting tRNA sequences are identical (38). The sequestered total tRNA^Phe^ was treated with RNase A to liberate the 3’-terminal adenosine and the reaction products were analyzed by LC-HRMS (**Figure 2D**). When extracted for the mass of Phe-Ade, the resulting ion chromatogram consists of two major peaks whose retention times and mass spectra match those detected *in vitro* and correspond to the 2’ and 3’-isomers of Phe-Ade (**Figure 2E and Supplementary Information, Figure S2**). When extracted for the mass of Ade, the ion chromatogram consists of a single major peak whose retention time also matches that of the *in vitro* sample (**Figure 2E and Supplementary Information, Figure S2**). Diacylation was not observed, and integration of the Phe-Ade and Ade EIC peaks indicate an *in vivo* acylation yield of 56% (39). No Phe-Ade was detected when the total RNA was extracted using o-Pyl, a biotin-tagged DNA oligonucleotide complementing tRNA^Pyl^ rather than tRNA^Phe^, or when RNase A was withheld (**Supplementary Information, Figure S3, Table S1**). These control experiments verify that RNase A can be used to assess the tRNA acylation level of material isolated from cells using biotin-tagged capture oligonucleotides (19).

### Controls and internal standard facilitates comparisons across PARTI samples

To compare acylated tRNA levels across multiple samples, we modified the PARTI analysis to include an internal standard, the peptide leucine-enkephalin (Leu-Enk) (40). To establish the improved precision provided by the internal Leu-Enk standard, we again isolated total RNA from *E. coli* and captured the tRNA^Phe^ population using o-Phe and streptavidin-coated magnetic beads as described. In this case, a fixed amount of Leu-Enk was added to each sample after RNase A cleavage and before LC-HRMS analysis. EIC traces for the expected masses of Phe-Ade and Leu-Enk show two Phe-Ade peaks, as expected, and one Leu-Enk peak with a retention time of 4.5 min (**Figure 3A**). The combined area of the Phe-Ade peaks was normalized to that of Leu-Enk to generate a unitless normalized value, A_norm_. We note that for established pulldown conditions using 250 µg total RNA, A_norm_ values were consistent between five biological replicates over two days, whereas the raw Phe-Ade peak areas varied by data collection day, indicating the benefit of the Leu-Enk internal standard (**Figure 3B, C**).

**Figure 3:**
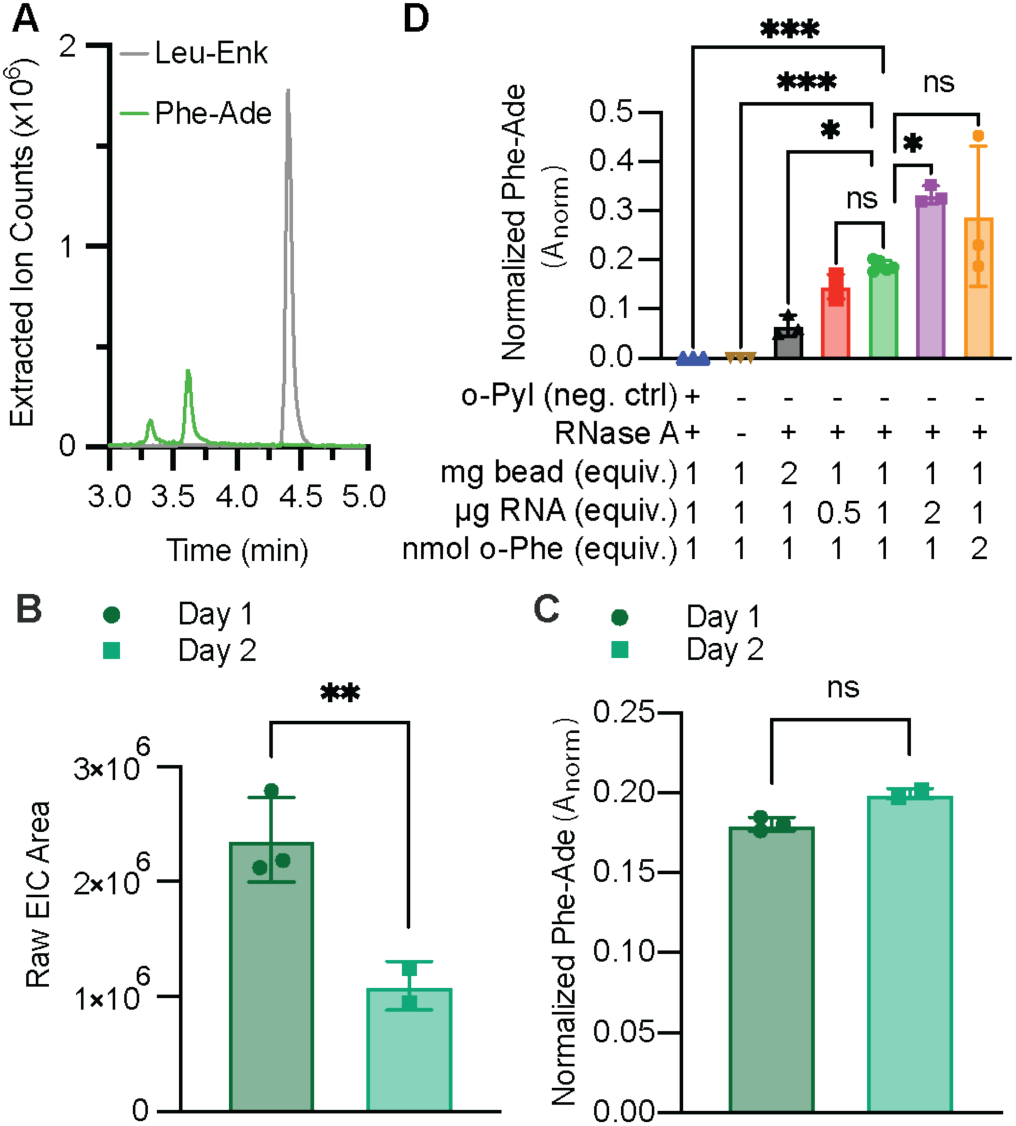
Controls and an internal standard facilitate comparisons across PARTI samples. (**A)** Shown are overlaid extracted ion chromatograms (EICs) of Leu-Enk (gray, calc [M+H]: 556.2766 Da) and Phe-Ade (green, calc [M+H]: 415.1724 Da). **(B)** Shown is a bar graph comparing raw Phe-Ade peak area sums collected by LC-HRMS over different days. Each point shown corresponds to a biological replicate and error bars represent one standard deviation from the average. Data acquired on Day 1 is shown in dark green (mean = 2.36 x 10^6^; SD = 3.68 x 10^6^; n = 3), whereas data acquired on Day 2 is shown in turquoise (mean = 1.10 x 10^6^; SD = 2.11 x 10^5^; n = 2). Statistical analysis bars represent the results of a two-tailed t-test. P > 0.05 = ns; p ≤ 0.05 = *; p ≤ 0.01 = **. **(C)** Shown is a bar graph comparing Phe-Ade peak areas normalized to Leu-Enk in each sample collected by LC-HRMS over different days. Samples shown are the same as in **(B).** Each point shown corresponds to a biological replicate and error bars represent one standard deviation from the average. Data from Day 1 is shown in dark green (mean = 0.18; SD = 0.00; n = 3), whereas data from Day 2 is shown in turquoise (mean = 0.20; SD = 0.00; n = 2). Statistical analysis bars represent the results of a two-tailed t-test. p > 0.05 = ns; p ≤ 0.05 = *; p ≤ 0.01 = **. **(D)** Shown is a bar graph displaying A_norm_ values for Phe-Ade detected using PARTI. Each point shown corresponds to a biological replicate and error bars represent one standard deviation from the average. Shown in green are A_norm_ values for Phe-Ade determined under standard conditions: 250 µg total RNA per sample, 5 mg streptavidin-coated magnetic beads, and 500 pmol o-Phe, as described in Methods (mean = 0.19; SD = 0.01; n = 5). Shown in red are A_norm_ values for Phe-Ade determined when total RNA per sample = 125 µg (mean = 0.15; SD = 0.02; n = 3). Shown in purple are A_norm_ values when total RNA per sample = 500 µg (mean = 0.33; SD = 0.02; n = 3). Shown in blue are A_norm_ values determined using 500 pmol o-Pyl (mean = 0; SD = 0; n = 3). Shown in orange is the A_norm_ value determined using 1 nmol o-Phe (mean = 0.29; SD = 0.14; n = 3). Shown in black is is the A_norm_ value determined when 1 mg (2x) streptavidin beads were used (mean = 0.07; SD = 0.02; n = 3). Statistical analysis bars represent the results of a two-tailed t-test. p > 0.05 = ns; p ≤ 0.05 = *; p ≤ 0.01 = **; p ≤ 0.001 = ***.

In addition to including an internal standard, we also performed experiments to evaluate the dependence of A_norm_ on multiple experimental parameters, including the concentrations of total RNA and the biotinylated DNA capture oligonucleotide as well as the mass of streptavidin-coated magnetic beads (**Figure 3D**). Compared to A_norm_ values obtained from the established conditions shown in green, doubling the streptavidin bead mass reduced the A_norm_ value by 63%, perhaps due to inefficient RNase A cleavage. Doubling the o-Phe concentration had no effect, providing evidence for near quantitative tRNA extraction (**Figure 3D**). We also observed that the A_norm_ value was 74% higher when the total RNA was doubled and 21% lower when the total RNA was halved relative to the standard value of 250 µg (**Figure 3D**). This finding implies that a total RNA mass of 250 µg does not saturate the volume of beads used. Capture using an oligomer complementary to tRNA^Pyl^ instead of o-Phe, or capture with o-Phe but without RNase A, yielded no Phe-Ade signal (**Figure 3D**).

### Validating o-Pyl as a capture oligonucleotide to sequester tRNA^Pyl^ from cells

With a validated PARTI workflow in place, we sought to apply it to evaluate the reactivity of an orthogonal tRNA/aaRS pair in cells supplemented with non-canonical α- and non-α-amino acids. We chose PylRS from *M. alvus*, which acylates *M. alvus* tRNA^Pyl^ with α-hydroxy acids and β^2^-hydroxy acids carrying appropriate side chains (7, 9, 41). To capture the complete tRNA^Pyl^ population, we designed a new biotinylated DNA capture oligonucleotide (o-Pyl) that is complementary to nucleotides 27 through 48 of tRNA^Pyl^ (**Figure 4A** and **Supplementary Information, Table S1**). To test the utility of o-Pyl, tRNA^Pyl^ was transcribed *in vitro*, purified, and acylated with L-α-boc-lysine (BocK) using purified *M. alvus* PylRS (**Figure 4A, B**). The tRNA^Pyl^ population was captured with o-Pyl, sequestered using streptavidin-coated magnetic beads, and treated with RNase A. The final sample generated using the PARTI protocol was doped with Leu-Enk and analyzed using LC-HRMS. Under these conditions, the yield of BocK-tRNA^Pyl^ is 37% in line with previous reports (7) (**Supplementary Information, Figure S4**). As expected, the EICs revealed the presence of Leu-Enk and BocK-Ade bound to the 2’ or 3’ end of adenosine (BocK-Ade) (**Figure 4B**). Only unreacted and monoacyl tRNA, but no diacyl tRNA, was detected, as observed previously (7) (**Supplementary Information, Figure S4**).

**Figure 4:**
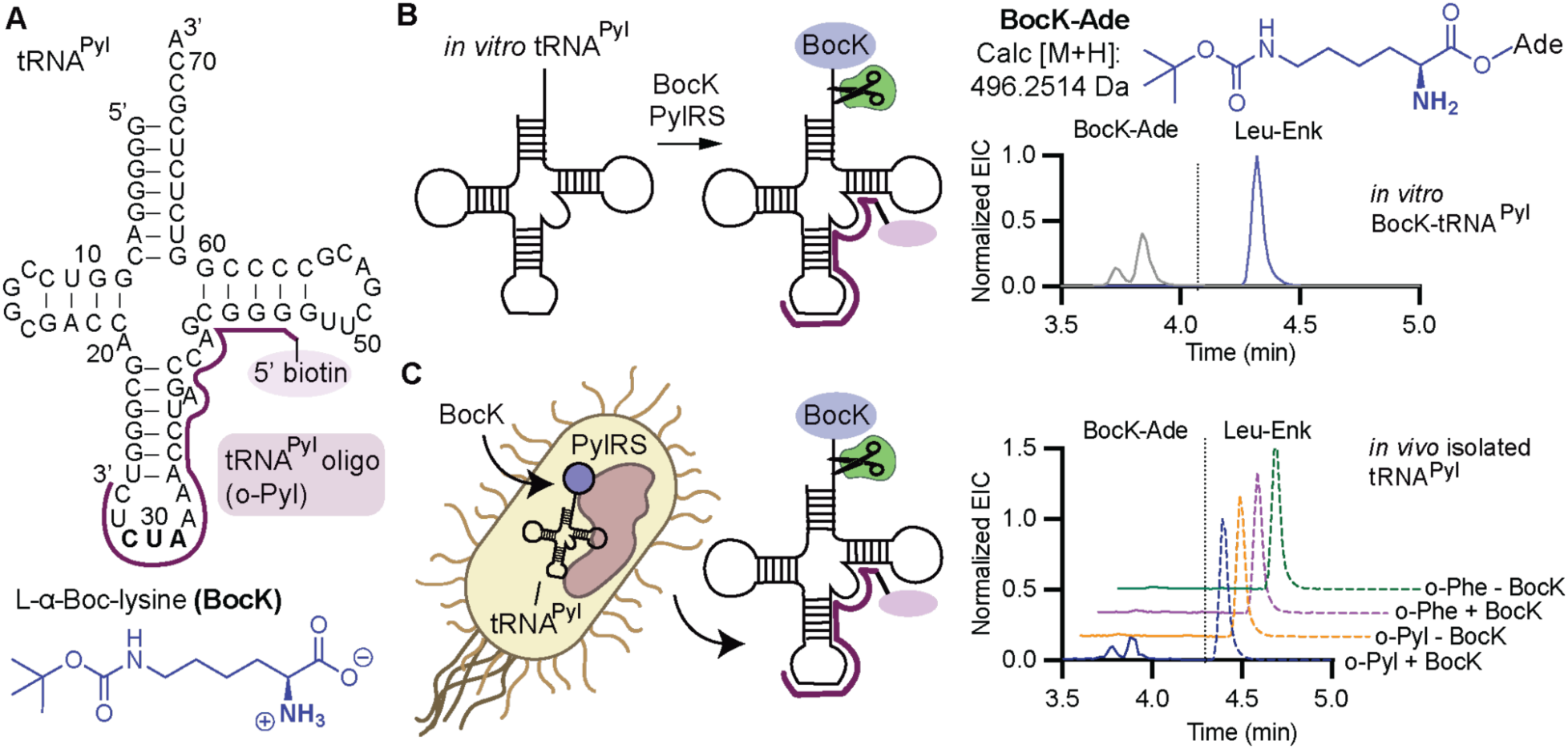
Validating PARTI *in vitro* and in cells using *M. alvus* PylRS/tRNA^Pyl^ and BocK. **(A)** The secondary structure of *M. alvus* tRNA^Pyl^ bound to o-Pyl and the structure of BocK. **(B)** *In vitro* acylation of tRNA^Pyl^ with PylRS and BocK followed by PARTI analysis produces the extracted ion chromatograms (EICs) shown at right. The trace in indigo represents the EIC for the mass of BocK-Ade; the trace in gray represents the EIC for the mass of Leu-Enk. **(C)** Acylation of tRNA^Pyl^ in cells followed by PARTI analysis produces the extracted ion chromatograms (EICs) shown at right. The four traces represent samples from *E. coli* DH5α cells expressing the *M. alvus* PylRS/tRNA^Pyl^ pair and grown with or without BocK and extracted with either o-Phe or o-Pyl. BocK-Ade (calc [M+H]: 496.2514 Da) is detected only in the presence of BocK and when the total RNA is extracted with o-Pyl. Each trace is normalized to Leu-Enk (dashed line, calc [M+H]: 556.2766 Da) in each sample.

Next, we sought to establish that A_norm_ values were dependent on the amount of sequestered tRNA^Pyl^ subjected to the PARTI workflow. To do so, we chose a monomer that results in detectable levels of both mono- and diacyl-tRNA^Pyl^ *in vitro* in the presence of PylRS – L-α-hydroxy-Boc-lysine (OH-BocK) (7, 9). *In vitro*-transcribed and purified tRNA^Pyl^ was acylated with OH-BocK in the presence of PylRS to yield a mixture of unreacted, monoacylated, and diacylated tRNA^Pyl^ (**Supplementary Information, Figure S5A-D**). Varying amounts of the mixed tRNA^Pyl^ population were sequestered with o-Pyl, treated with RNase A, and analyzed by LC-HRMS (**Supplementary Information, Figure S5E**). We found that A_norm_ values for detection of either acylated adenosine species trend linearly with the amount of starting tRNA^Pyl^ (**Supplementary Information, Figure S5F**).

As a final control, we grew *E. coli* expressing the *M. alvus* PylRS/tRNA^Pyl^ pair in the presence or absence of the PylRS substrate BocK. We extracted total RNA, sequestered the tRNA^Pyl^ population with o-Pyl and streptavidin-coated magnetic beads, treated the beads with RNase A, and analyzed the products using LC-HRMS (**Figure 4C**). Total RNA was also extracted with o-Phe. When BocK and o-Pyl were both present, BocK-Ade was detected alongside Leu-Enk in the EIC at the same retention time as in the *in vitro* control (**Figure 4 B, C**). In contrast, when BocK was present but o-Phe was used, or when BocK was absent and either o-Pyl or o-Phe were used, the BocK-Ade signal was absent (**Figure 4C**).

### PARTI reveals a difference in cellular acylation of tRNA^Pyl^ with *(R)*- and *(S)*-β_2_-hydroxy acids

Next, we applied PARTI to evaluate the acylation state of tRNA^Pyl^ in cells expressing PylRS and supplemented with either *(R)-* or *(S)*-β_2_-hydroxy-boc-lysine (β_2_-OH-BocK) (**Figure 5A**).

**Figure 5:**
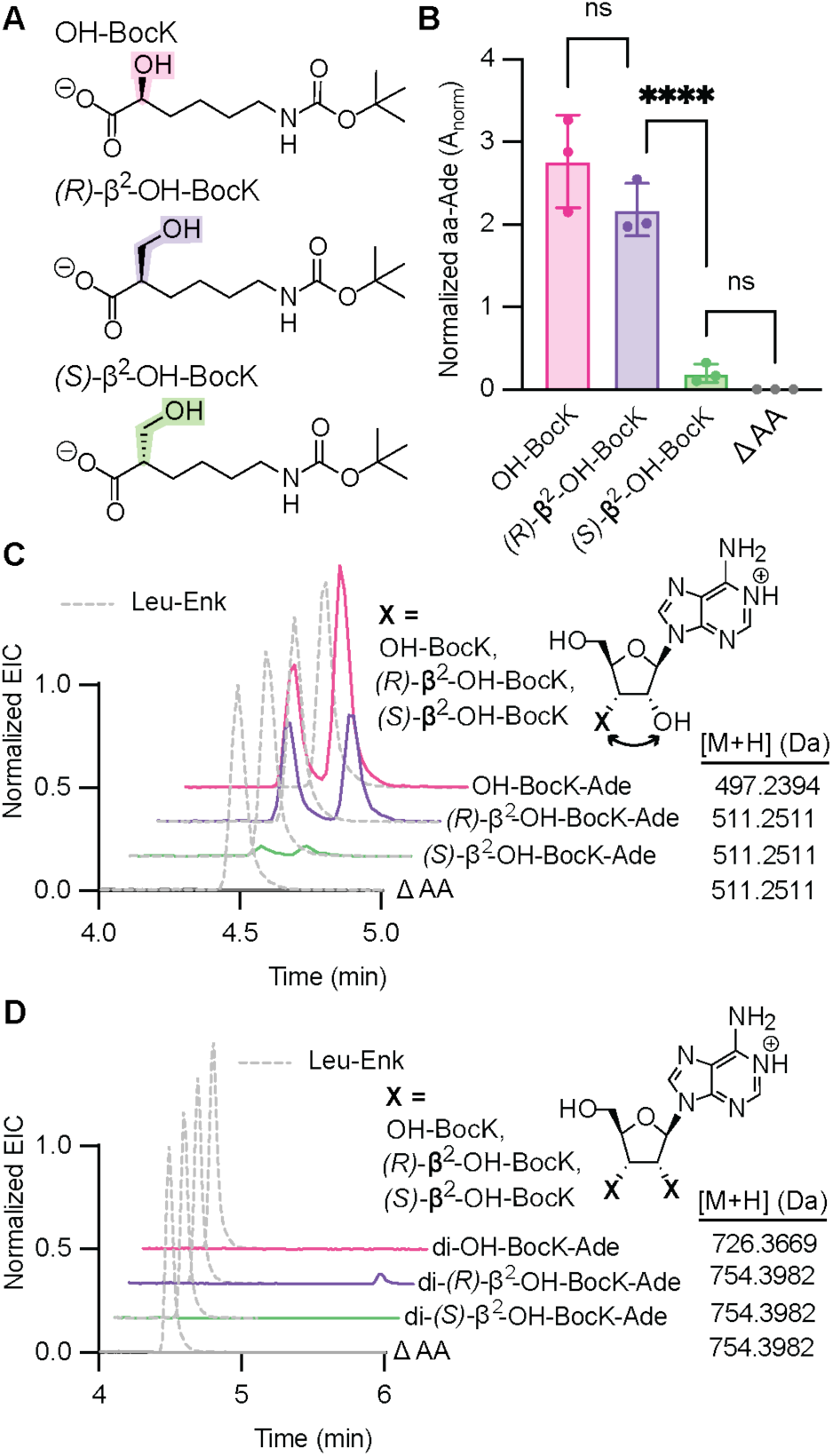
*(R)-* and *(S)-*β^2^-hydroxy acids are not equivalent substrates for PylRS in cells. **(A)** Structures of OH-BocK, (*R)*-β^2^-OH-BocK, and (*S)*-β^2^-OH-BocK. (**B)** Shown is a bar graph displaying the relative A_norm_ values when 0.1 mM of each monomer in **(A)** is added to *E. coli* C321 cells expressing the *Ma*PylRS/tRNA^Pyl^ and PylRS and subjected to the PARTI workflow. Points shown correspond to biological replicates. Error bars represent one standard deviation from the average. Data representing the detection of OH-BocK-Ade is shown in pink (mean = 2.76; SD = 0.56; n = 3), detection of (*R)*-β_2_-OH-BocK-Ade is shown in purple (mean = 2.18; SD = 0.32; n = 3) and detection of (*S)*-β_2_-OH-BocK-Ade is shown in green (mean = 0.20; SD = 0.11; and n = 3). No β_2_-OH-BocK-Ade was observed when no substrate was added, as shown in gray (mean = 0.0; SD = 0.0; n = 3). Statistical analysis bars represent the results of a one-way ANOVA. p > 0.05 = ns; p ≤ 0.05 = *; p ≤ 0.01 = **; p ≤ 0.001 = ***. **(C)** Overlaid extracted ion chromatograms (EICs) of monoacyl-3’ Ade species detected by LC-HRMS following PARTI with *E. coli* C321 cells expressing the *Ma*PylRS/tRNA^Pyl^ pair and grown with no substrate or 0.1 mM OH-BocK, (*R)*-β_2_-OH-BocK, or (*S)*-β_2_-OH-BocK. Traces are normalized to the Leu-Enk EIC in each sample, shown in dashed gray. **(D)** Overlaid extracted ion chromatograms (EICs) of diacylated 3’ Ade species detected by LC-MS following PARTI with *E. coli* C321 cells expressing tRNA^Pyl^ and PylRS grown with no substrate or 0.1 mM OH-BocK, (*R)*-β_2_-OH-BocK or (*S)*-β_2_-OH-BocK. Traces are normalized to the Leu-Enk EIC in each sample, shown in dashed gray.

Previous work has shown that both β_2_-OH-BocK stereoisomers are substrates for the *M. alvus* PylRS/tRNA^Pyl^ pair *in vitro*, but only one stereoisomer–*(R)-*β_2_-OH-BocK–is incorporated into proteins in cells (7). In addition, although both β_2_-OH-BocK enantiomers are processed by PylRS *in vitro* as well as OH-BocK, under comparable conditions the yield of protein containing *(R)-*β_2_-OH-BocK was approximately ten-fold lower than the yield of protein containing OH-BocK (7). We used PARTI to determine whether the level of tRNA^Pyl^ acylation by these monomers in cells could account for either or both of these observations.

We grew *E. coli* C321 cells expressing the *Ma*PylRS/tRNA^Pyl^ pair in the presence of 0.1 mM *(R)-* or *(S)*-β_2_-OH-BocK or OH-BocK (**Figure 5A**), extracted total RNA, and isolated the tRNA^Pyl^ population using o-Pyl and streptavidin-coated magnetic beads. The beads were treated with RNase A and the eluted products characterized using LC-HRMS (**Figure 5B, C and Supplementary Information, Figure S6, S7**). Under these expression conditions, which are identical to those used for cellular experiments reported previously (7), the extent of tRNA^Pyl^ acylation depends on both β_2_-OH-BocK stereochemistry and backbone identity (α- or β) (**Supplementary Information, Figure S8**). When the growths were supplemented with (*R)*-β_2_-OH-BocK, the A_norm_ value resulting from monoacylation of tRNA^Pyl^ by (*R)*-β^2^-OH-BocK-Ade was comparable to that of OH-BocK-Ade (**Figure 5B**), in line with the reported yields of these two acylated tRNAs *in vitro* (7). However, when the growths were supplemented with (*S)*-β_2_-OH-BocK, the A_norm_ value for (*S)*-β^2^-OH-BocK-Ade was approximately 10–fold lower than that for either OH-BocK-Ade or *(R)*-β_2_-OH-BocK-Ade (**Figure 5B)**. These observations indicate that in cells, only a relatively small fraction of the available tRNA^Pyl^ is acylated with (*S)*-β_2_-OH-BocK. The first implication of this finding is that the previously reported selectivity for incorporation of *(R)*-β_2_-OH-BocK in cells is due at least in part to differences in tRNA^Pyl^ acylation. Either *(S)*-β_2_-OH-BocK is a poor PylRS substrate in cells, or it is metabolized into an unknown product that is no longer a PylRS substrate. The second implication is that the low level of incorporation of *(R)*-β_2_-OH-BocK in cells relative to OH-BocK is due to bottlenecks that occur after tRNA^Pyl^ acylation. One likely bottleneck is EF-Tu, which engages poorly *in vitro* with tRNA^Phe^ when it is acylated with *(R)*-or *(S)*-β_2_-Phe or, notably, with *(R)*-or *(S)*-β_3_-Phe (11), but other bottlenecks cannot be ruled out.

Another interesting feature of the acylation of tRNA^Pyl^ with *(R)-* or *(S)-*β_2_-OH-BocK *in vitro* was the appearance of diacylated tRNA products (7). Diacylated tRNAs–tRNAs acylated on both the 2’ and 3’-hydroxyl groups–were first observed decades ago when *T. thermophilus* PheRS was used to acylate *E. coli* tRNA^Phe^ with Phe *in vitro* (12). Several years later it was reported that a modified variant of *E. coli* tRNA^Ala^ chemically diacylated with alanine or allylglycine supported the incorporation of these monomers into protein in an *E. coli* cell lysate (13). More recent work detected various diacylated tRNA^Pyl^ species *in vitro* when tRNA^Pyl^ was reacted with PylRS or variants thereof in the presence of OH-BocK, *(R)-* and *(S)*-β_2_-OH-BocK, des-amino BocK, and L-α-amino-, α-thio-, and L-α-hydroxy-Phe (7, 9). As far as we know, diacylated tRNAs have never been detected in cells.

We applied the PARTI workflow to evaluate whether tRNA^Pyl^ is diacylated in cells expressing PylRS in the presence of OH-BocK, *(R)-*β_2_-OH-BocK, or *(S)-*β_2_-OH-BocK. No evidence for tRNA diacylation was observed when the cells were supplemented with either OH-BocK or *(S)*-β_2_-OH-BocK (**Figure 5D** and **Supplementary Information, Figure S6, S7**). Diacylation of tRNA^Pyl^ was detected, however, in cells supplemented with *(R)*-β_2_-OH-BocK (**Figure 5D** and **Supplementary Information, Figure S6, S7**). The di-*(R)*-β_2_-OH-BocK-Ade detected from RNA isolated from cells was identical in both elution time and exact mass to di-*(R)*-β_2_-OH-BocK-Ade detected after *in vitro* tRNA^Pyl^ acylation reactions (**Supplementary Information, Figure S6, S7, S9**). *In vitro*, the relative levels of mono- and diacylated tRNA^Pyl^ are related linearly to enzyme concentration (7). The relative levels of mono- and diacylated tRNA^Pyl^ detected from cells correspond to *in vitro* acylation reactions performed previously which used less than 2.5 µM PylRS, 10 mM *(R)*-β_2_-OH-BocK, and 25 µM tRNA^Pyl^ (7) (**Supplementary Information, Figure S5**). Although the detected level of diacylated tRNA^Pyl^ in cells is relatively low, we cannot rule out that it does not contribute, at least in part, to the lower yield of intact protein containing *(R)*-β_2_-OH-BocK.

### PARTI detects *in vivo* acylation of monomers that have not yet been reported as ribosome elongation substrates

There is great interest in expanding ribosome chemistry beyond simple amine and hydroxyl/thiol nucleophiles to generate ketone products containing a CC bond in place of the canonical CN or CO/CS bonds. Ribosome products containing backbone ketones can be generated by post-translational acyl rearrangements (5), but have not yet been detected as direct ribosome products. Direct CC bond formation within the ribosome active site demands one or more aaRS enzymes that acylate tRNA with a monomer capable of establishing a proximal carbon-centered nucleophile under physiological conditions. Previous work has shown that PylRS and several variants thereof (FRS1, FRS2, FRSA) acylate tRNA^Pyl^ *in vitro* with benzylmalonate derivatives capable of generating a carbon-centered nucleophile after decarboxylation, as occurs during reactions catalyzed by polyketide synthases (9, 42, 43). Benzylmalonate derivatives shown to be substrates for the PylRS variant FRSA include *meta*-trifluoromethyl-2-benzylmalonate (*m*-CF_3_-bma), *meta*-bromo-2-benzylmalonate (*m*-Br-bma), and *meta*-methyl-2-benzylmalonate (*m*-CH_3_-bma) (**Figure 6A**) (9). We used PARTI to establish whether FRSA could acylate tRNA^Pyl^ in cells with each of these benzylmalonate derivatives.

**Figure 6:**
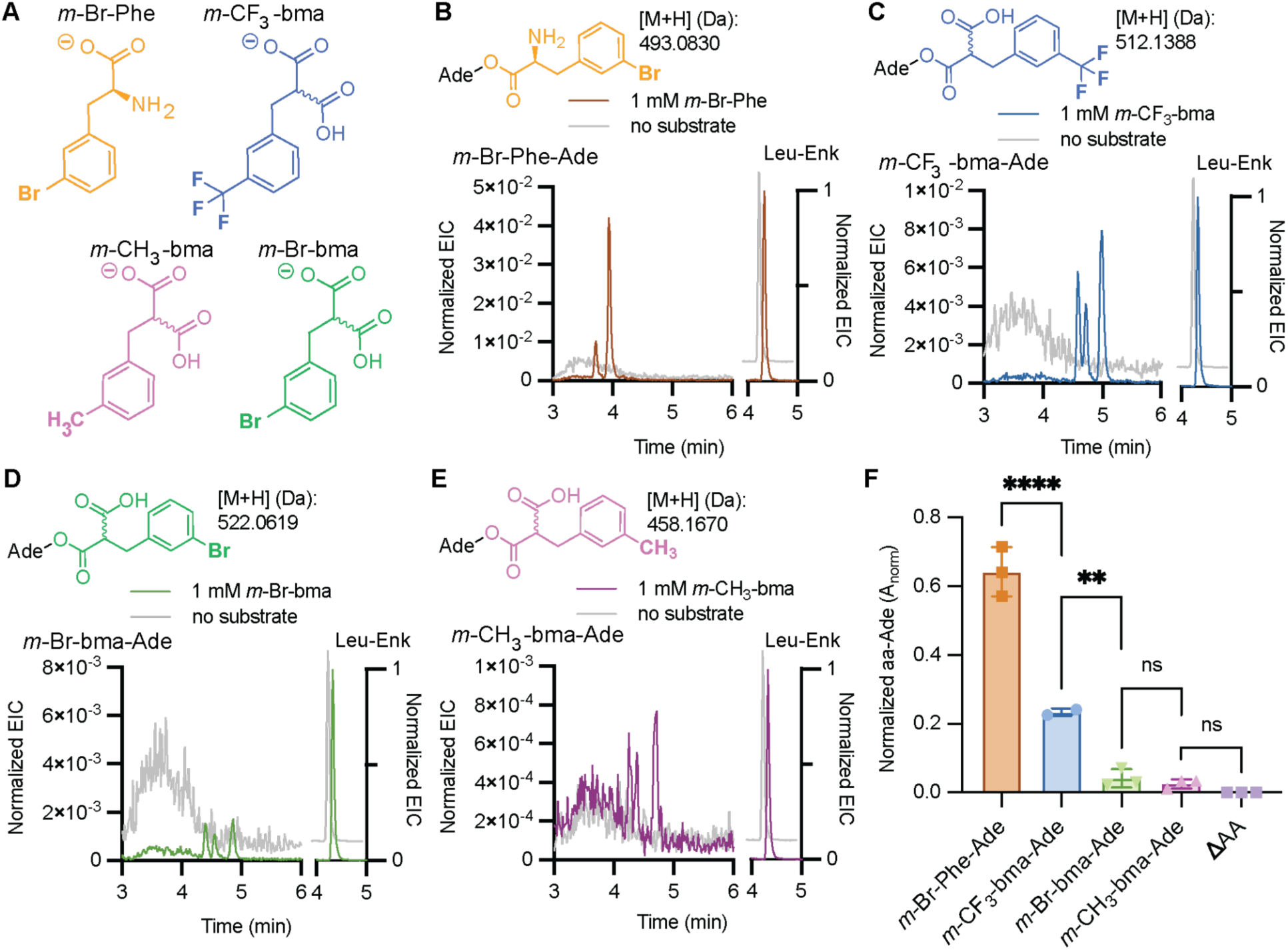
Benzylmalonate derivatives are substrates for FRSA in cells. **(A)** Structures of *meta*-bromo-phenylalanine (*m*-Br-Phe), *meta*-bromo-2-benzylmalonate (*m*-Br-bma), *meta*-methyl-2-benzylmalonate (*m*-CH_3_-bma) and *meta*-trifluoromethyl-2-benzylmalonate (*m*-CF_3_-bma). **(B)** Structure and calculated mass of *m-*Br-Phe-Ade and overlaid EICs of calculated [M+H] for *m-*Br-Phe-Ade, (at left, [M+H]: 493.0830 Da) normalized to Leu-Enk (at right, [M+H]: 556.2766 Da) after PARTI using RNA isolated from *E. coli* C321 cells expressing *Ma*FRSA and *Ma*tRNA^Pyl^ and grown with (brown) or without (gray) 1 mM *m-*Br-Phe. **(C)** Structure and calculated mass of *m-*CF_3_-bma-Ade and overlaid EICs of calculated [M+H] for *m-* CF_3_-bma-Ade, (at left, [M+H]: 512.1388 Da) normalized to Leu-Enk (at right, [M+H]: 556.2766 Da) after PARTI with RNA isolated from *E. coli* C321 cells expressing *Ma*FRSA and *Ma*tRNA^Pyl^ and grown with (blue) or without (gray) 1 mM *m-*CF_3_-bma. **(D)** Structure and calculated mass of *m-*Br-bma-Ade and overlaid EICs of calculated [M+H] for *m-*Br-bma-Ade, (at left, [M+H]: 522.0619 Da) normalized to Leu-Enk (at right, [M+H]: 556.2766 Da) after PARTI with RNA from *E. coli* C321 cells expressing *Ma*FRSA and *Ma*tRNA^Pyl^ and grown with (green) or without (gray) 1 mM *m-*Br-bma. **(E)** Structure and calculated mass of *m-*CH_3_-bma-Ade and overlaid EICs of calculated [M+H] for *m-*CH_3_-bma-Ade, (at left, [M+H]: 458.1670 Da) and Leu-Enk (at right, [M+H]: 556.2766 Da) after PARTI with RNA from *E. coli* C321 cells expressing *Ma*FRSA and *Ma*tRNA^Pyl^ and grown with (pink) or without (gray) 1 mM *m-*CH_3_-bma. **(F)** Shown is a bar graph displaying relative amounts of monoacyl-Ade recovered from *E. coli* C321 cells grown with 1 mM of the corresponding benzylmalonate or α-amino acid substrate and expressing tRNA^Pyl^ and FRSA. Points shown correspond to biological replicates and error bars represent one standard deviation from the average. Data representing the detection of *m*-Br-Phe-Ade is shown in orange (mean = 0.64; SD = 0.07; n = 3). Data representing the detection of *m*-CF_3_-bma, *m*-Br-bma, and *m*-CH_3_-bma are shown in blue (mean = 0.23; SD = 0.01; n = 2), green (mean = 0.04; SD = 0.03; n = 3), and pink (mean = 0.03; SD = 0.01; n= 3), respectively. Statistical analysis bars represent the results of a one-way ANOVA. p > 0.05 =ns; p ≤ 0.05 = *; p ≤ 0.01 = **; p ≤ 0.001 = ***.

To this end, total RNA was isolated from *E. coli* C321 cells expressing the *Ma*FRSA/tRNA^Pyl^ pair and supplemented with 1 mM of either *meta*-bromo-phenylalanine (*m*-Br-Phe) as a positive control or one of the three benzylmalonate substrates, and the tRNA^Pyl^ isolated and analyzed via the standard PARTI workflow. In the case of *m*-Br-Phe as a substrate, extracting the total ion chromatogram for the mass of *m*-Br-Phe-Ade generates an EIC with two isobaric peaks which were not observed when the cells were grown in the absence of added substrate (**Figure 6B**). In the case of the three benzylmalonate substrates, three peaks were present in each of the EICs which were absent when substrate was not added during cell growth (**Figure 6C-E, Supplementary Information, Figure S10**). As many as four isobaric peaks are expected when tRNA is monoacylated with a pro-chiral benzylmalonate derivative, as the reaction generates the expected mixture of 2’ and 3’ products that is each a mixture of two diastereomers (9). Multiple isobaric peaks in the EIC were also observed when tRNA^Pyl^ was acylated *in vitro* with these substrates and FRSA (9).

Evaluation of the A_norm_ values resulting from acylation of tRNA^Pyl^ in cells with three benzylmalonate substrates shows a pattern of reactivity that parallels the yields of acylated tRNA^Pyl^ observed *in vitro* (9). These A_norm_ values imply that the most reactive benzyl malonate in cells is *m*-CF_3_-bma, followed by *m*-Br-bma and then *m*-CH_3_-bma; the ratio of A_norm_ values for these three substrates in cells was 8:3:1 (**Figure 6F**). These A_norm_ values are 3-, 15-, and 25-fold lower than that observed for the non-canonical α-amino acid which served as a positive control, *m-*Br-Phe (**Figure 6F**) (42). Notably, although the relative cellular activities of benzylmalonate substrates implied by A_norm_ values parallels acyl-tRNA^Pyl^ yields observed *in vitro* (9), the substrate-dependent differences are greater in cells. *In vitro*, using 10 mM benzylmalonate, 25 µM tRNA^Pyl^ and 2.5 µM FRSA, the detected ratio of 3-adenylated products was 3:2:1 for *m*-CF_3_-bma, *m*-Br-bma and *m*-CH_3_-bma, respectively (9). Thus *m*-CF_3_-bma is a substantially better substrate in cells than expected based on *in vitro* reactivity.

### Evidence for cellular metabolism of *N*-Me BocK

There is also great interest in the cellular biosynthesis of *N*-methylated proteins and peptides, as this modification can alter protein conformation and improve stability, bioavailability, target affinity, and selectivity (44, 45). Amide *N*-Me groups can be installed within peptides prepared synthetically using solid phase methods, or using enzymes, or using the ribosome and chemically acylated tRNAs *in vitro* and in cell lysate (23, 28, 30, 46–49). However, *N*-methylated proteins have not yet been prepared in live cells for reasons thought to be associated with poor accommodation of *N*-Me aminoacyl-tRNA by the ribosome (22, 25) and/or low affinity for elongation factor Tu (EF-Tu) (29). Several aminoacyl synthetases have been reported to acylate tRNA with *N*-Me amino acids *in vitro*. For example, *N*-methyl BocK (*N*-Me BocK) is a substrate for *Methanosarcina mazei* PylRS (41), and *N*-methylated phenylalanine analogs are substrates for *M. alvus* PylRS variants (9).

In alignment with these examples, we found that *N*-Me BocK is also a substrate for *Ma*PylRS *in vitro*. Incubation of 5 µM *M. alvus* PylRS with 25 µM tRNA^Pyl^ and 10 mM BocK resulted in 29% BocK-tRNA^Pyl^ as determined using intact tRNA LC-MS; when the analogous reaction was performed with *N*-Me BocK, the yield was 26% (**Supplementary Information, Figure S11**).

Even so, when *E. coli* Top10 or C321 cells expressing *Ma*PylRS, *Ma*tRNA^Pyl^ and a superfolder GFP (sfGFP) plasmid containing a single TAG codon at position 200 (sfGFP-200TAG) were supplemented with *N*-Me BocK, the only GFP product isolated contained BocK, not *N*-Me BocK (**Figure 7A** and **Supplementary Information, S12**). The yield of purified sfGFP-200TAG containing BocK when cells were supplemented with *N*-Me BocK (8 mg/L) was more than 20% of the yield isolated when cells were supplemented with the equivalent concentration of BocK (36 mg/L). There was no evidence for BocK contamination in the NMR or high-resolution mass spectrum of the *N*-Me BocK stock (**Supplementary Information, Figure S13A, B**), nor was BocK-Ade detected from LC-HRMS analysis following RNase A treatment of tRNA^Pyl^ acylated *in vitro* with *Ma*PylRS and *N*-Me BocK (**Supplementary Information, Figure S14A**).

**Figure 7:**
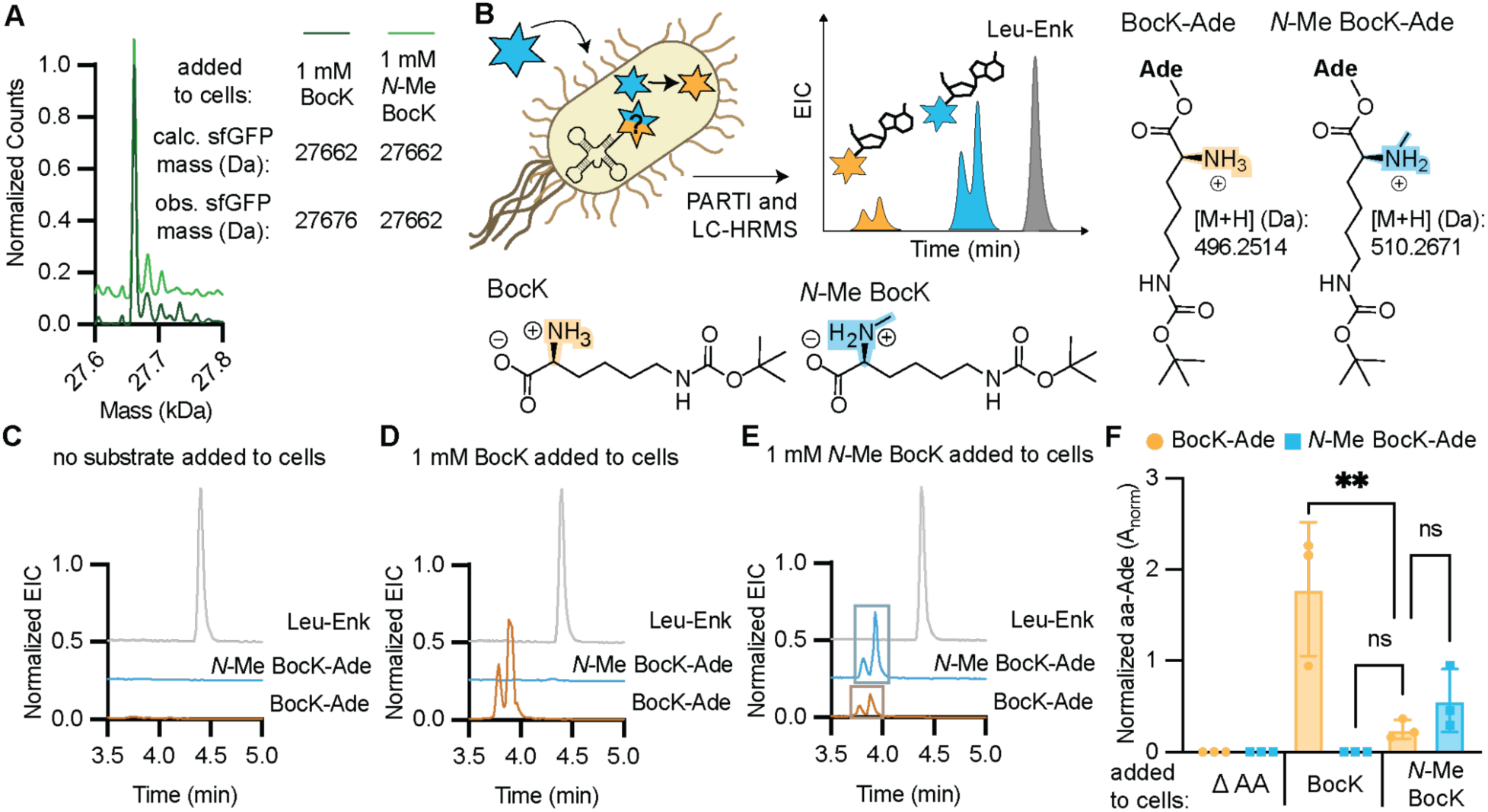
Metabolism of PylRS Substrates *in vivo* is observable from PARTI with tRNA^Pyl^. **(A)** Deconvoluted LC-HRMS spectra of sfGFP-200TAG purified from *E. coli* Top10 cells supplemented with 1 mM BocK (light green, expected mass: 27,662.3 Da) or 1 mM N-Me BocK (dark green, expected mass: 27,676.3 Da). The observed mass of both proteins is 27,662.3 Da. Signal is normalized to the highest count value within each sample and the trace illustrating the sfGFP-200TAG mass when the growth were supplemented with *N*-Me BocK is shifted upwards on the y-axis for visibility. **(B)** Schematic summarizing the PARTI workflow for observing populations of acylation of tRNA^Pyl^ with *N*-Me BocK (blue) as well as its metabolic product BocK (yellow). Also shown are the structures of the RNase A cleavage products BocK-Ade (expected monoacyl mass: 496.2514 Da) or *N*-Me BocK-Ade (expected monoacyl mass: 510.26707). **(C)** Overlaid EICs for detection of Leu-Enk (in gray, [M+H]: 556.2766 Da), BocK-Ade (in gold, [M+H]: 496.2514 Da), and *N*-Me BocK-Ade (in blue, [M+H]: 510.2671 Da) in a PARTI sample from *E. coli* Top10 cells expressing *Ma*PylRS and *Ma*tRNA^Pyl^ grown with no added substrate. **(D)** Overlaid EICs for detection of Leu-Enk (in gray, [M+H]: 556.2766 Da), BocK-Ade (in gold, [M+H]: 496.2514 Da), and *N*-Me BocK-Ade (in blue, [M+H]: 510.2671 Da) in a PARTI sample from *E. coli* Top10 cells expressing *Ma*PylRS and *Ma*tRNA^Pyl^ grown with 1 mM BocK. **(E)** Overlaid EICs for detection of Leu-Enk (in gray, [M+H]: 556.2766 Da), BocK-Ade (in gold and boxed in muted gold, [M+H]: 496.2514 Da), and *N*-Me BocK-Ade (in blue and boxed in muted blue, [M+H]: 510.2671 Da) in a PARTI sample from *E. coli* Top10 cells expressing *Ma*PylRS and *Ma*tRNA^Pyl^ grown with 1 mM *N*-Me BocK. (**F)** Shown is a bar graph displaying relative amounts of BocK-Ade (yellow) and *N*-Me BocK-Ade (blue) recovered from *E. coli* Top10 cells expressing tRNA^Pyl^ and PylRS and grown with 1 mM indicated monomer(s). PARTI was carried out using o-Pyl and graphed values are respective aa-Ade signals normalized to Leu-Enk signal within each LC-HRMS sample. Points in each bar correspond to biological replicates, and different species detected within a given cell growth condition are from the same biological samples. Error bars are one standard deviation from the average. When cells were supplemented with only 1 mM BocK, only BocK-Ade was detected (mean = 1.79 SD= 0.73 n=3). In cells supplemented with only 1 mM *N*-Me BocK, both *N-*Me-BocK Ade (mean = 0.57 SD= 0.34 n=3) and BocK-Ade (mean = 0.25 SD = 0.11 n= 3) were observed. When no substrate was added, neither BocK-Ade nor OH-BocK-Ade were detected. Statistical analysis bars represent the results of a one-way ANOVA. p > 0.05 =ns; p ≤ 0.05 = *; p ≤ 0.01 = **; p ≤ 0.001 = ***.

Regardless, to eliminate the possibility that BocK contamination in the *N*-Me BocK stock could account for the expression of sfGFP-200TAG containing BocK, we grew Top10 and C321 cells expressing *Ma*PylRS, *Ma*tRNA^Pyl^ and sfGFP-200TAG in the presence of between 1 pM and 1 mM BocK to determine both the lowest BocK concentration that would yield a detectable level of sfGFP-200TAG containing BocK and the required concentration to replicate the signal observed when 1 mM *N-*Me BocK had been added (**Supplementary Information, Figure S13C, D**). These results indicate that 1 nM BocK is the lowest BocK concentration that would yield a detectable level of sfGFP-200TAG containing BocK. Furthermore, even a 10% impurity of BocK in the N-Me BocK sample would be insufficient to support the level of incorporation observed in either Top10 or C321 *E. coli*, and the N-Me BocK monomer is at least 99.9% pure (**Supplementary Information, Figure S13E, F**). As BocK contamination in the *N*-Me BocK stock cannot account for the observed expression of sfGFP-200TAG containing BocK, we hypothesized that *N*-Me BocK was metabolized into BocK in cells at one or more points prior to translation.

To test this hypothesis, we used the PARTI workflow to probe the acylation state of tRNA^Pyl^ in *E. coli* expressing the PylRS/tRNA^Pyl^ pair and supplemented with *N*-Me BocK, and probed specifically for whether tRNA^Pyl^ was acylated with BocK, *N*-Me BocK, or a mixture of the two (**Figure 7B**). We incubated *E. coli* Top10 cells expressing the *Ma*PylRS/tRNA^Pyl^ pair with either 1 mM BocK, 1 mM *N*-Me BocK, or no substrate, and then subjected each isolated RNA population to the PARTI workflow. No peaks corresponding to the mass of either BocK-Ade or *N*-Me BocK-Ade were detected when no substrate was used to supplement the cell growths (**Figure 7C and Supplementary Information, Figure S14**). When cells were supplemented with only BocK, LC-HRMS revealed the anticipated pair of isomeric BocK-Ade peaks (**Figure 7D**). We verified that BocK-Ade captured from an *in vivo* sample exhibited the same retention times as the products of an *in vitro* tRNA^Pyl^ acylation reaction (**Supplementary Information, Figure S14**). No peaks corresponding to the mass of *N*-Me BocK-Ade were detected from either the *in vitro* or the *in vivo* sample (**Figure 7D and Supplementary Information, Figure S14**).

However, when we used the PARTI workflow to analyze total RNA isolated from cells supplemented with only *N*-Me BocK, we detected clear evidence of both *N*-Me BocK-Ade and BocK-Ade (**Figure 7E**). *N*-Me BocK-Ade was detected from the *in vivo* sample as two isobaric peaks that eluted at the same retention time as the products of the *in vitro* tRNA^Pyl^ acylation reaction (**Supplementary Information, Figure S14)** and were absent when no substrate was added to the cell growths (**Figure 7C and Supplementary Information, Figure S14).** BocK-Ade was also detected in the sample isolated from cells supplemented with only *N*-Me BocK as two peaks with a distinct elution time from *N*-Me BocK-Ade (**Figure 7E).** Again, BocK-Ade eluted at the same retention time as the products of an *in vitro* tRNA^Pyl^ acylation reaction (**Supplementary Information, Figure S14)**; these peaks were absent when no substrate was added to the cell growths (**Figure 7C and Supplementary Information, Figure S14**). The A_norm_ value for BocK-Ade in this sample was 86% lower than from the sample supplemented with 1 mM BocK and was approximately one-half the A_norm_ value due to *N*-Me-BocK-Ade (**Figure 7F**).

We carried out additional LC-HRMS experiments with known amounts of acylated *in vitro* tRNA^Pyl^ to assess if there were significant differences in the ionization efficiencies of BocK-Ade and *N*-Me-BocK-Ade. *In vitro-*purified and transcribed tRNA^Pyl^ was acylated with either BocK or *N-*Me BocK and the yields of BocK-tRNA^Pyl^ and *N-*Me BocK-tRNA^Pyl^, respectively, were determined using intact tRNA LC-MS (**Supplementary Information, Figure S11**). Aliquots from each reaction were treated with RNase A, doped with Leu-Enk, and analyzed using LC-HRMS to determine A_norm_ values for BocK-Ade or *N-*Me BocK-Ade. Each A_norm_ value was then divided by the yield of the acylation reaction. We observed that A_norm_/% acylation was 50% higher when BocK was the substrate (**Supplementary Information, Figure S15**), suggesting that BocK-Ade ionizes 50% more efficiently than *N*-Me BocK-Ade. When accounting for this difference, BocK-Ade levels are still only about four times lower than *N-*Me BocK-Ade levels when cells were supplemented with only *N*-Me BocK. The level of BocK contamination could not account for the observed level of BocK-Ade, supporting the hypothesis that *N*-Me-BocK is metabolized into BocK. Because no *N*-Me BocK is incorporated into protein, BocK appears to entirely outcompete *N*-Me BocK in the translation steps following acylation.

## DISCUSSION

Here we describe PARTI, a mass spectrometry-based assay that provides a snapshot of the cellular acylation state of a user-defined tRNA. PARTI differs from previously reported cellular tRNA acylation assays in terms of the information it provides. Unlike assays that rely on acyl-tRNA hydrolysis and tRNA amplification (15, 16, 18, 50–52), PARTI provides the exact mass of the acylating species rather than simply whether or not the tRNA has been acylated. In most cases, this exact mass is sufficient to confirm that acylation has occurred with the monomer of interest and not a metabolized derivative or a native α-amino acid. And, unlike assays that rely on acyl-tRNA hydrolysis and mass-guided monomer identification (16, 8, 19), PARTI differentiates between mono- and diacylated tRNA products and thereby does not over-estimate the state of tRNA acylation. Although billions of years of evolution have mitigated diacylation as a concern for native α-amino acids, this side reaction remains a concern for non-α-amino acid monomers, as it is unknown how these unusual tRNA species affect the sophisticated interplay of factors and the ribosome that embody rapid translation.

We applied the PARTI workflow here to more deeply scrutinize the multiple steps required to introduce non-α-amino acid monomers into proteins in cells. We focused first on β^2^-hydroxy acid monomers, which can be introduced into protein at two separate positions using the PylRS/tRNA^Pyl^ pair from *M. alvus* (7). Recent results show that although both enantiomers of β^2^-hydroxy-boc-lysine (β^2^-OH-BocK) are substrates for *M. alvus* PylRS *in vitro*, only the *(R)* enantiomer is introduced into protein in cells, and with yields 10-fold lower than anticipated based on *in vitro* tRNA acylation data (7). Computational results imply that enantioselectivity does not involve the ribosome directly (7). Using PARTI, we discovered that enantioselectivity is due at least in part to low steady-state levels of tRNA^Pyl^ acylated with *(S)*-β^2^-OH-BocK but not *(R)*-β^2^-OH-BocK. *(S)*-β^2^-OH-BocK may fail to enter cells, be altered by metabolization once it arrives, or it may fail to generate a stable acylated tRNA^Pyl^ upon reaction with PylRS.

We also discovered using PARTI that the lower level of incorporation of *(R)*-β^2^-OH-BocK relative to OH-BocK in cells is likely due to factors that follow tRNA acylation, as the steady state levels of the two acylated tRNAs are comparable. Indeed, recent work has shown that both *(R)* and *(S)* enantiomers of β^2^- and β^3^-Phe disrupt ternary complex formation with EF-Tu by at least an order of magnitude *in vitro* and the complexes that do form ostensibly bypass the proofreading stage of mRNA decoding (11). This finding emphasizes that for monomers that are not α-amino acids, myriad other translation factors as well as the ribosome itself (7) must also be considered before heteropolymers containing multiple copies of these unusual substrates, alone or in combination, can be prepared at scale. We note that PARTI was also used here to verify that three different benzyl malonate derivatives are sufficiently cell-permeant to support robust levels of tRNA acylation. This finding demonstrates how PARTI may be used as a screen to establish the relative activities of monomers that are not yet ribosome substrates.

The final discovery facilitated by PARTI in this work relates to *N*-Me-α-amino acids that are of enormous current interest for the development of peptide-derived therapeutics *N*-Me-α-amino acids can be introduced into peptides in reconstituted *in vitro* translation mixtures (22–29, 53) and in S-30 cell extracts (30), and some are excellent substrates for aaRS enzymes *in vitro* (9, 41). Yet there are no examples in which even a single *N*-Me-α-amino acid has been introduced into a protein in a cell. PARTI revealed that when *E. coli* expressing the PylRS/tRNA^Pyl^ pair is supplemented with *N*-Me-BocK, a notable fraction of the isolated tRNA^Pyl^ carried BocK in addition to the portion acylated with *N*-Me-BocK. Furthermore, incorporation of only BocK into sfGFP was observed, implying that *N*-Me-BocK is metabolized *via* an unknown pathway into BocK in cells. Further work will be required to establish whether this metabolism targets the free amino acid or the acyl-tRNA and whether other *N*-alkyl amino acids are equally affected.

Indeed, it is likely that the efficient incorporation of *N*-Me backbones *in vivo* may require strain engineering, as used previously to mitigate the metabolic conversion of α-hydroxy acids into α-amino acids (3, 9).

In summary, the PARTI assay reported here effectively bridges the informational gap between *in vitro* acylation and cellular translation. Its ease and simplicity should benefit ongoing efforts to study and improve the cellular incorporation of non-α-amino acid monomers into proteins.

## DATA AVAILABILITY

All data in the manuscript are available in the Supplementary Data, and raw data is available upon request.

## ACKNOWLEDGEMENTS

The authors are grateful to members of the Schepartz labs for helpful discussion, and especially to Lauren Lesiak, Angel Vázquez Maldonado, and Dr. Daniel Brauer for comments on the manuscript. We also thank Leah Roe for providing benzylmalonate substrates, Noah Hamlish for chemically competent C321 cells, and Alex Solivan for obtaining NMR data.

