## Supplementary Information for "Monitoring monomer-specific acyl-tRNA levels in cells with PARTI"

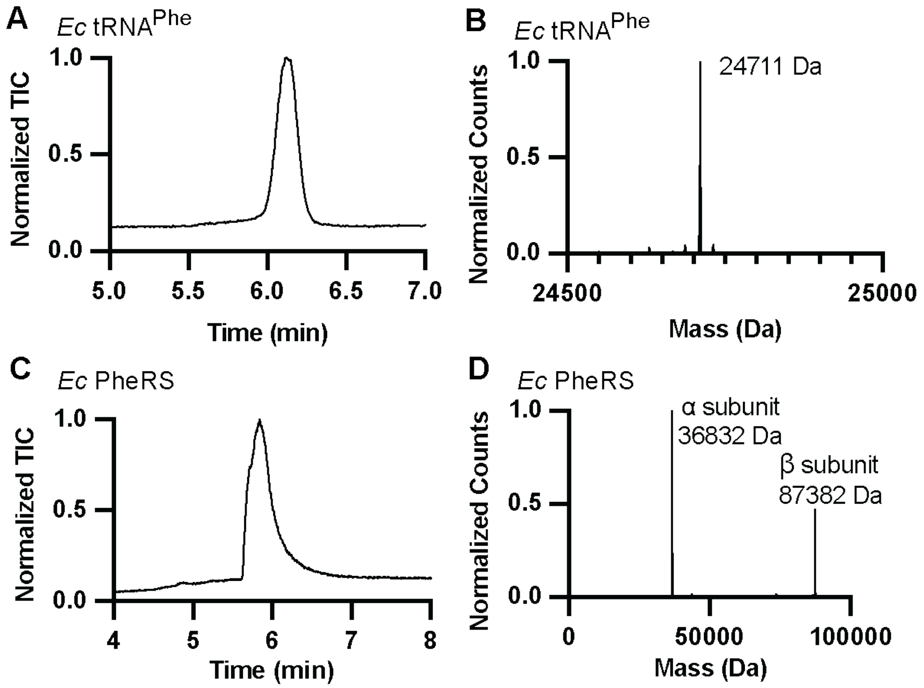

**Figure S1: Purified *Ec*PheRS and *Ec*tRNA$^{Phe}$ characterized by LC-MS.** Shown are the **(A)** total ion chromatogram (TIC) and **(B)** deconvoluted mass spectra for *in vitro*-transcribed and purified *Ec*tRNA$^{Phe}$ (expected mass: 24,710 Da). Shown are the **(C)** TIC and **(D)** deconvoluted mass spectrum for purified *Ec*PheRS (expected α subunit mass: 36830.82 Da, expected β subunit mass: 87378.11 Da).

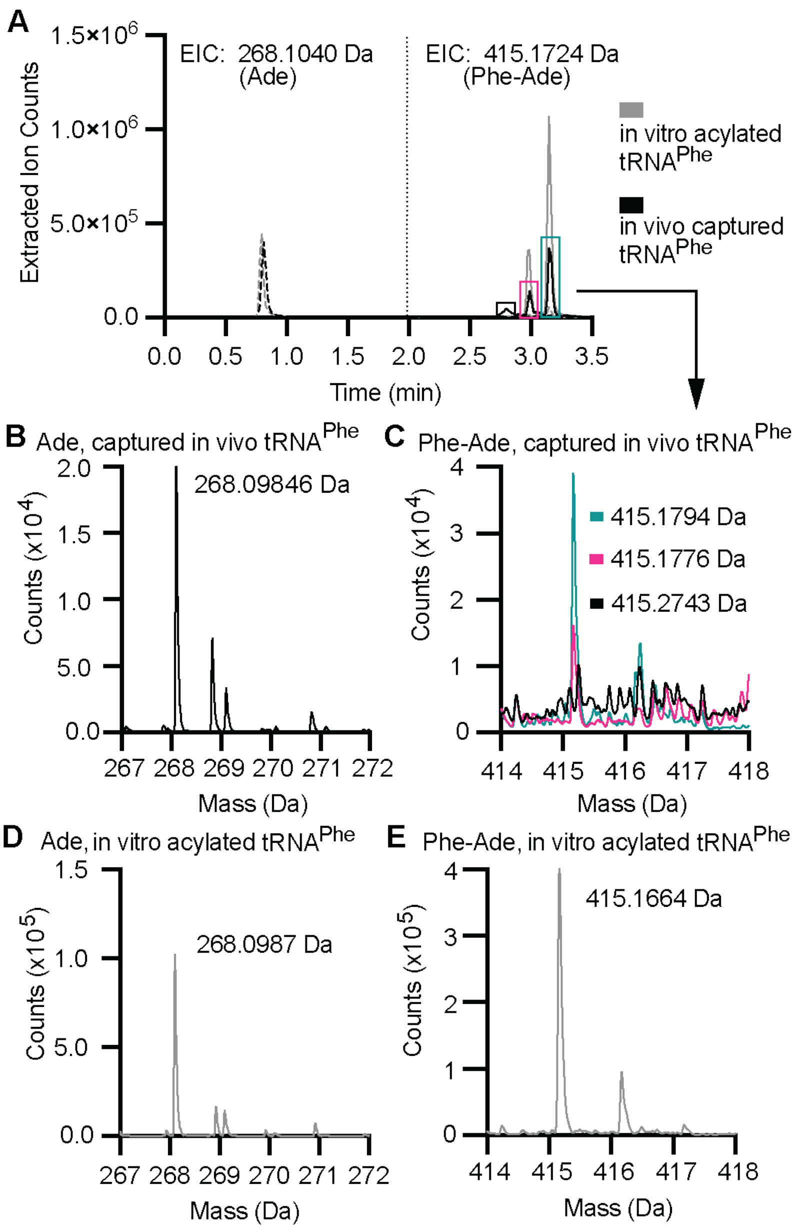

**Figure S2: Acylation of tRNA$^{Phe}$ in cells using PARTI generates the same products as those detected *in vitro*.** Experimental details are identical to those in **Figure 2**. **(A)** Shown are overlaid extracted ion chromatograms (EICs) of Ade (dotted line, calc [M+H]): 268.1040 Da) and Phe-Ade (solid line, calc [M+H]): 415.1724 Da) detected following RNase A treatment of tRNA$^{Phe}$ acylated *in vitro* with Phe and PheRS (gray) or from cells after capture with o-Phe (black). The peaks from the *in vivo* sample that align with the *in vitro* sample are boxed in pink and teal while the small peak exclusively observed *in vivo* is boxed in black. **(B)** The mass spectrum of Ade detected following isolation and RNase A cleavage of *in vivo* tRNA$^{Phe}$. **(C)** Shown are overlaid mass spectra extracted from each EIC peak of the *in vivo* Phe-tRNA$^{Phe}$ sample boxed in **(A)**. The pink and teal traces (obs: 415.1794 Da and 415.1776 Da) were counted towards Phe-Ade yield. The black peak (415.2743 Da) was excluded due to its presence in blank samples. **(D)** The mass spectrum of Ade detected in the cleaved *in vitro* Phe-tRNA$^{Phe}$ sample is shown. **(E)** The mass spectrum of Phe-Ade detected in the cleaved *in vivo* Phe-tRNA$^{Phe}$ purified with PARTI is shown.

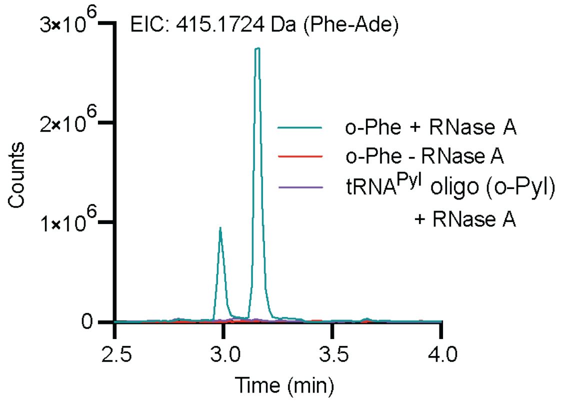

**Figure S3: Phe-Ade is not detected from purified *E. coli* tRNA when RNase A is withheld or when a non-complementary DNA capture oligonucleotide is used**. Shown are EIC traces from LC-HRMS analysis of three PARTI samples where RNA from *E. coli* DH5α cells was processed with either o-Phe and RNase A (teal), o-Phe but no RNase A (red) or o-Pyl and RNase A (purple) as detailed in Methods. Traces shown are EICs for the expected mass of Phe-Ade (calc [M+H]: 415.1724 Da).

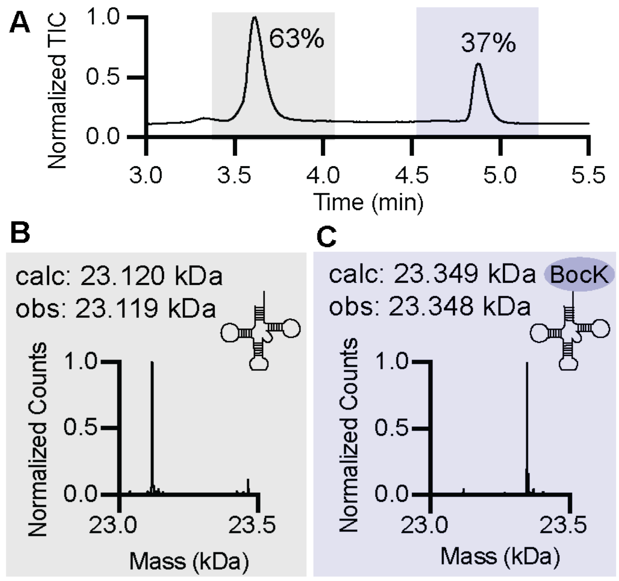

**Figure S4: Intact tRNA LC-MS of the products of *in vitro* tRNA$^{Pyl}$ acylation confirms presence of unreacted tRNA$^{Pyl}$ and BocK-tRNA$^{Pyl}$.** Shown is the TIC of tRNA$^{Pyl}$ purified from an aminoacylation reaction containing 25 µM tRNA$^{Pyl}$, 10 µM *M. alvus* PylRS, and 10 mM BocK incubated for 2 h at 37°C. The peak highlighted in gray corresponds to unreacted tRNA$^{Pyl}$ and the peak in purple corresponds to BocK-tRNA$^{Pyl}$. Deconvoluted mass spectra of **(B)** unreacted tRNA$^{Pyl}$ or **(C)** BocK-tRNA$^{Pyl}$ derived from the highlighted peaks in **(A).** The respective expected and observed masses of each product are shown.

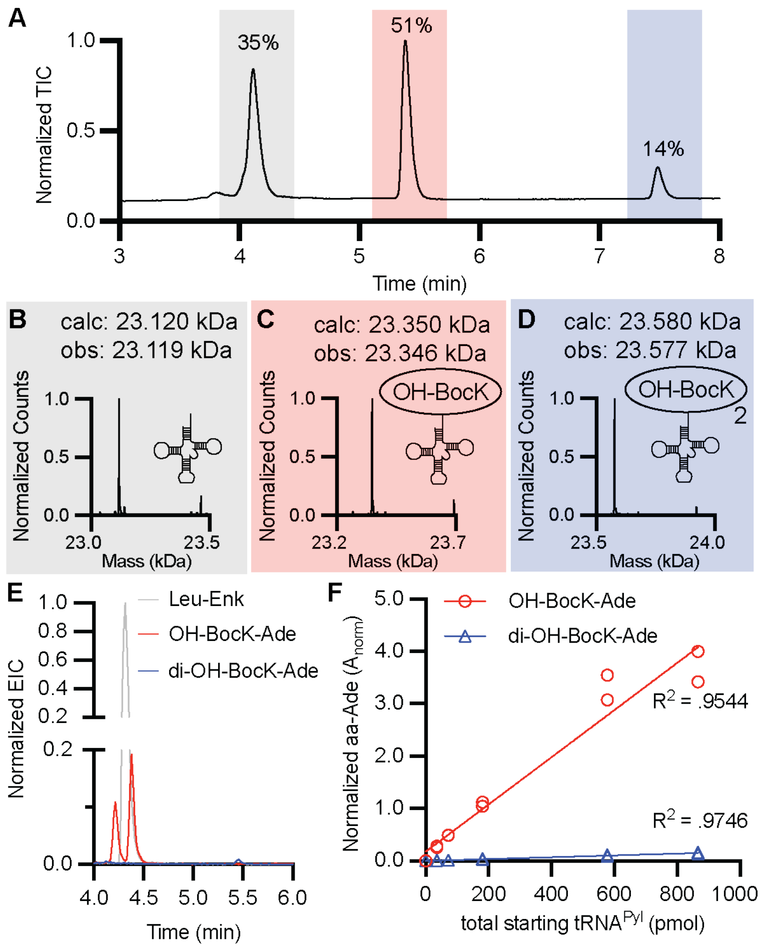

**Figure S5: $A_{norm}$ values for OH-BocK-Ade and di-OH-BocK-Ade correlate linearly with the amount of *in vitro* tRNA$^{Pyl}$ processed with PARTI. (A)** Shown is the TIC recorded during intact tRNA LC-MS of an *in vitro* acylation of 25 µM tRNA$^{Pyl}$ with 10 mM OH-BocK and 10 µM purified *M. alvus* PylRS after 2 h at 37 °C. The highlighted TIC peaks correspond to unreacted tRNA$^{Pyl}$ (gray), OH-BocK-tRNA$^{Pyl}$ (red), and di-OH-BocK-tRNA$^{Py}$ (blue), respectively. **(B-D)** The corresponding deconvoluted mass spectra for each highlighted TIC peak are shown with a comparison between their expected and observed mass. **(E)** Overlaid EICs of calculated [M+H] for OH-BocK-Ade (red, [M+H]: 497.2394 Da) and di-OH-BocK-Ade (blue, [M+H]: 726.3669 Da) normalized to Leu-Enk (gray, [M+H]: 556.2766 Da) following PARTI with the sample characterized in **(A-D). (F)** Shown is a plot of $A_{norm}$ values from OH-BocK-Ade (red) and di-OH-BocK-Ade (blue) recorded by LC-HRMS from PARTI reactions using varying amounts of the tRNA$^{Pyl}$ acylation reaction characterized in **(A-D)**. The amount of total tRNA$^{Pyl}$ was determined as described using a NanoDrop ND-1000 device**.** Then, a dilution series of the reaction was made in technical duplicate and PARTI was performed on each dilution. $A_{norm}$ values from OH-BocK-Ade and di-OH-BocK-Ade were determined for each replicate and plotted against the amount of total tRNA$^{Pyl}$ in each PARTI experiment.

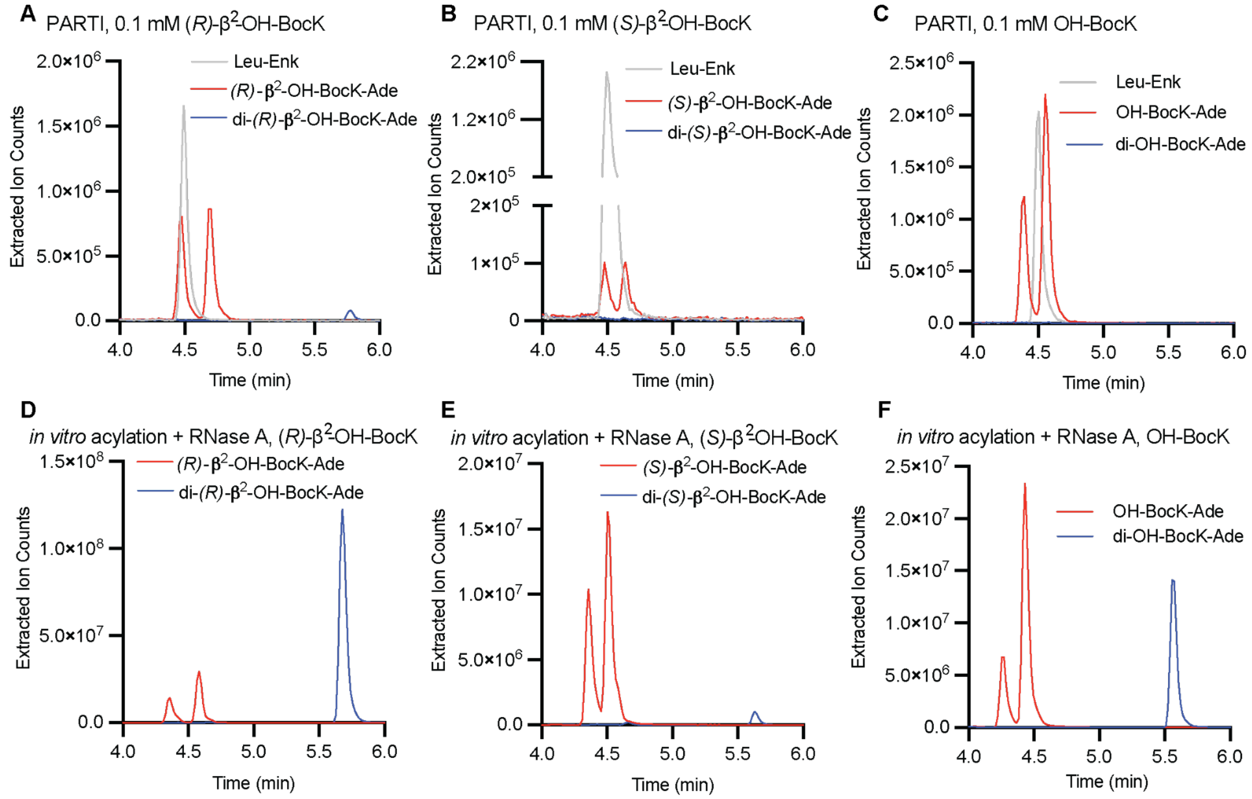

**Figure S6: LC-HRMS peaks for mono- and di-acylation of hydroxy acids charged to tRNA$^{Pyl}$ *in vitro* enable confirmation of *in vivo* acylation after PARTI.** PARTI was performed on RNA from *E. coli* C321 cells expressing *Ma*PylRS and *Ma*tRNA$^{Pyl}$ and grown with 0.1 mM of: **(A)** (*R*)-β$^2$-OH-BocK **(B)** (*S*)-β$^2$-OH-BocK or **(C)** OH-BocK. Overlaid EICs of Leu-Enk standard (gray), monoacylated (red) and diacylated (blue) products of each reaction are shown. Overlaid EICs of Leu-Enk standard (gray), monoacylated (red) and diacylated (blue) products of each reaction are shown. In parallel, *in vitro* tRNA$^{Pyl}$ acylations were carried out for 2 h at 37°C with 12.5 µM PylRS when **(D)** (*R*)-β$^2$-OH-BocK and **(E)** (*S*)-β$^2$-OH-BocK were the substrate and with 2.5 µM PylRS when **(F)** OH-BocK was the substrate. Substrate concentrations were 10 mM and the tRNA$^{Pyl}$ concentration was 25 µM. From each reaction 250 pmol total tRNA$^{Pyl}$ were treated with the RNase A assay as described and the equivalent of 10 pmol cleaved tRNA$^{Pyl}$ were analyzed by LC-HRMS as described in Methods. Overlaid EICs of monoacylated (red) and diacylated (blue) products of each reaction are shown. Calculated [M+H] values: Leu-Enk = 556.2766 Da, mono-(*R*)-β$^2$-OH-BocK-Ade = 511.2511 di-(*R*)-β$^2$-OH-BocK-Ade = 754.3982 Da, mono-(*S*)-β$^2$-OH-BocK-Ade = 511.2511 Da, di-(*S*)-β$^2$-OH-BocK-Ade = 754.3982 Da, mono-OH-BocK-Ade = 497.2394 Da and di-OH-BocK-Ade = 726.3669 Da.

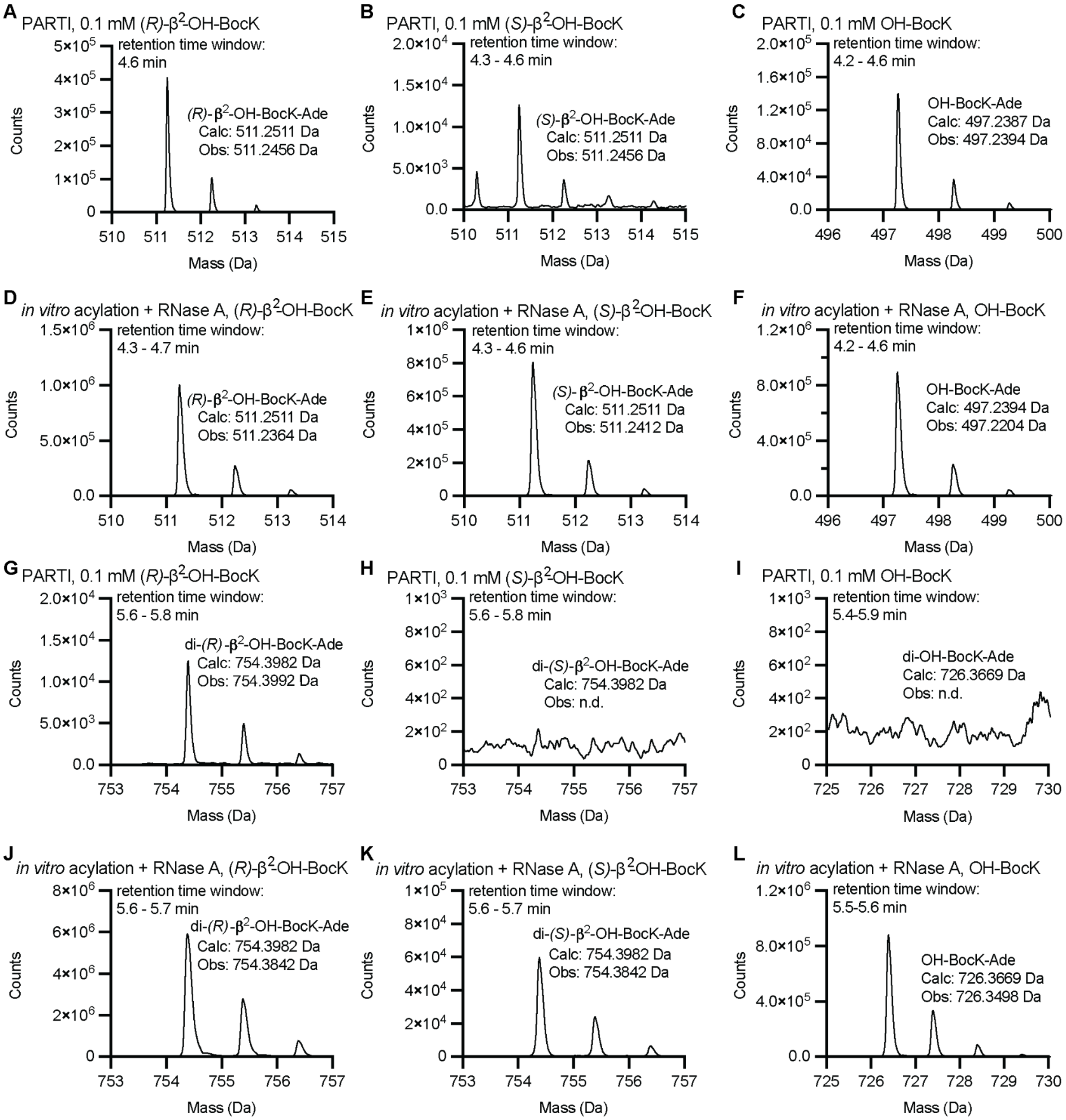

**Figure S7:** Mass spectra of aa-Ade detected *in vitro* and *in vivo* correspond to expected masses. All calculated masses are [M+H], and all spectra are extracted from samples in Supplementary Figure 6 at the indicated elution times. Shown are mass spectra of **(A)** (*R)*-β$^2$-OH-BocK-Ade, **(B)** (*S)*-β$^2$-OH-BocK-Ade, and **(C)** OH-BocK-Ade detected after PARTI was performed on RNA from *E. coli* C321 cells expressing *Ma*PylRS and *Ma*tRNA$^{Pyl}$ and grown with 0.1 mM of (*R)*-β$^2$-OH-BocK, (*S)*-β$^2$-OH-BocK, or OH-BocK, respectively. Shown are mass spectra of **(D)** (*R)*-β$^2$-OH-BocK-Ade, **(E)** (*S)*-β$^2$-OH-BocK-Ade, and **(F)** OH-BocK-Ade detected following RNase A treatment of *in vitro* tRNA$^{Pyl}$ acylated with (*R)*-β$^2$-OH-BocK, (*S)*-β$^2$-OH-BocK, or OH-BocK, respectively. Shown are mass spectra of **(G)** di-(*R)*-β$^2$-OH-BocK-Ade, **(H)** di-(*S)*-β$^2$-OH-BocK-Ade (not detected), and **(I)** OH-BocK-Ade (not detected) after PARTI was performed on RNA from *E. coli* C321 cells expressing *Ma*PylRS and *Ma*tRNA$^{Pyl}$ and grown with 0.1 mM of (*R)*-β$^2$-OH-BocK, (*S)*-β$^2$-OH-BocK, or OH-BocK, respectively. Shown are mass spectra of **(J)** di-(*R)*-β$^2$-OH-BocK-Ade, **(K)** di-(*S)*-β$^2$-OH-BocK-Ade, and **(L)** di-OH-BocK-Ade detected following RNase A treatment of *in vitro* tRNA$^{Pyl}$ acylated with (*R)*-β$^2$-OH-BocK, (*S)*-β$^2$-OH-BocK, or OH-BocK, respectively.

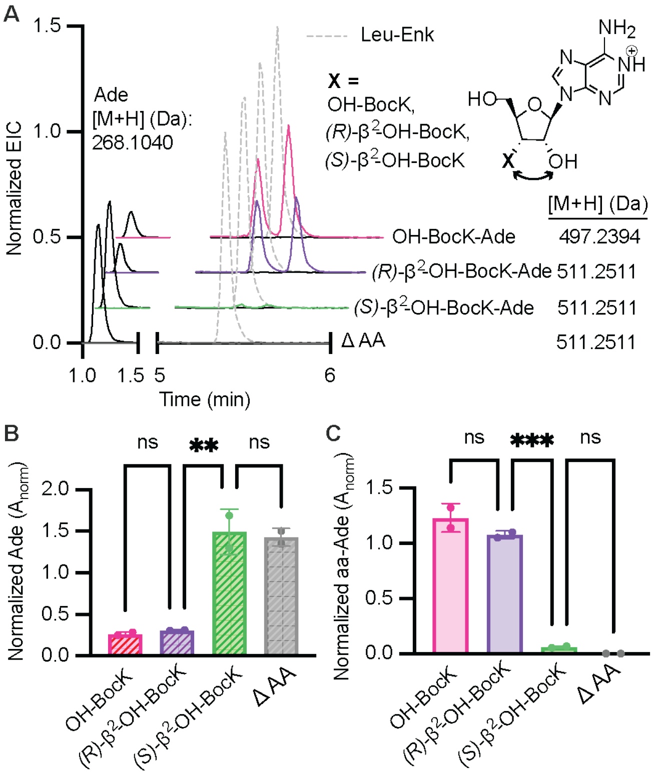

**Figure S8**: Monoacylation relates inversely to free 3' adenosine captured with PARTI. **(A)** Overlaid EICs of free 3' adenosine (in black) and monoacylated species detected by LC-HRMS following PARTI with *E. coli* C321 cells expressing $tRNA^{Pyl}$ and PylRS grown with no substrate or 0.1 mM OH-BocK (pink), (*R*)-$\beta^2$-OH-BocK (purple), or (*S*)-$\beta^2$-OH-BocK (green). Traces are normalized to the Leu-Enk EIC in each sample, shown in dashed gray. **(B)** Shown is a bar graph displaying relative amounts of unreacted 3' Ade recovered from *E. coli* C321 cells grown with 0.1 mM respective monomer and expressing $tRNA^{Pyl}$ and PylRS. PARTI was carried out using o-Pyl and graphed values are respective Ade signals normalized to the Leu-Enk signal within each LC-HRMS sample. Ade detected from cells grown with 0.1 mM OH-BocK (mean = 0.27 SD = 0.02), (*R*)-$\beta^2$-OH-BocK (mean = 0.31 SD = 0.00), (*S*)-$\beta^2$-OH-BocK (mean = 1.49 SD = 0.28), and no added substrate (mean = 1.43 SD = 0.11) are shown. Experiments were carried out as described in Methods except with changes to the chromatography protocol and MS collection window during LC-HRMS. Mobile phase B was initially held at 2% for 1 minute followed by a linear gradient from 2 to 4% over 1.89 minutes. Then, mobile phase B underwent a gradient from 4 to 40% over 3.11 minutes and a gradient from 40 to 100% over 2 minutes. Mobile phase B then transitioned from 100 to 4% over 2 minutes then was held at 4% for 0.5 minutes. Mass spectrometry data was collected between 0.72 and 8 min. **(C)** Shown is a bar graph displaying relative amounts of monoacyl monomers recovered from the same *E. coli* C321 cells as in **(B)** and graphed values are respective aa-Ade signals normalized to Leu-Enk signal within each LC-HRMS sample. Shown is OH-BocK-Ade (calc [M+H]: 497.2394 Da, mean = 1.23 SD = 0.13), (*R*)-$\beta^2$-OH-BocK-Ade (calc [M+H]: 511.2511 Da, mean = 1.07 SD = 0.03) and (*S*)-$\beta^2$-OH-BocK-Ade (calc [M+H]: 511.2511 Da, mean = 0.06 SD = 0.01). No $\beta^2$-OH-BocK-Ade (calc [M+H]: 511.2511 Da) was observed when no substrate was added (mean = 0.0 SD = 0.0). For all bar graphs each point corresponds to a biological replicate (n = 2) and error bars represent one standard deviation from the average. Statistical analysis bars represent the results of a one-way ANOVA. $p > 0.05$ =ns; $p \leq 0.05$ = *; $p \leq 0.01$ = **; $p \leq 0.001$ = ***.

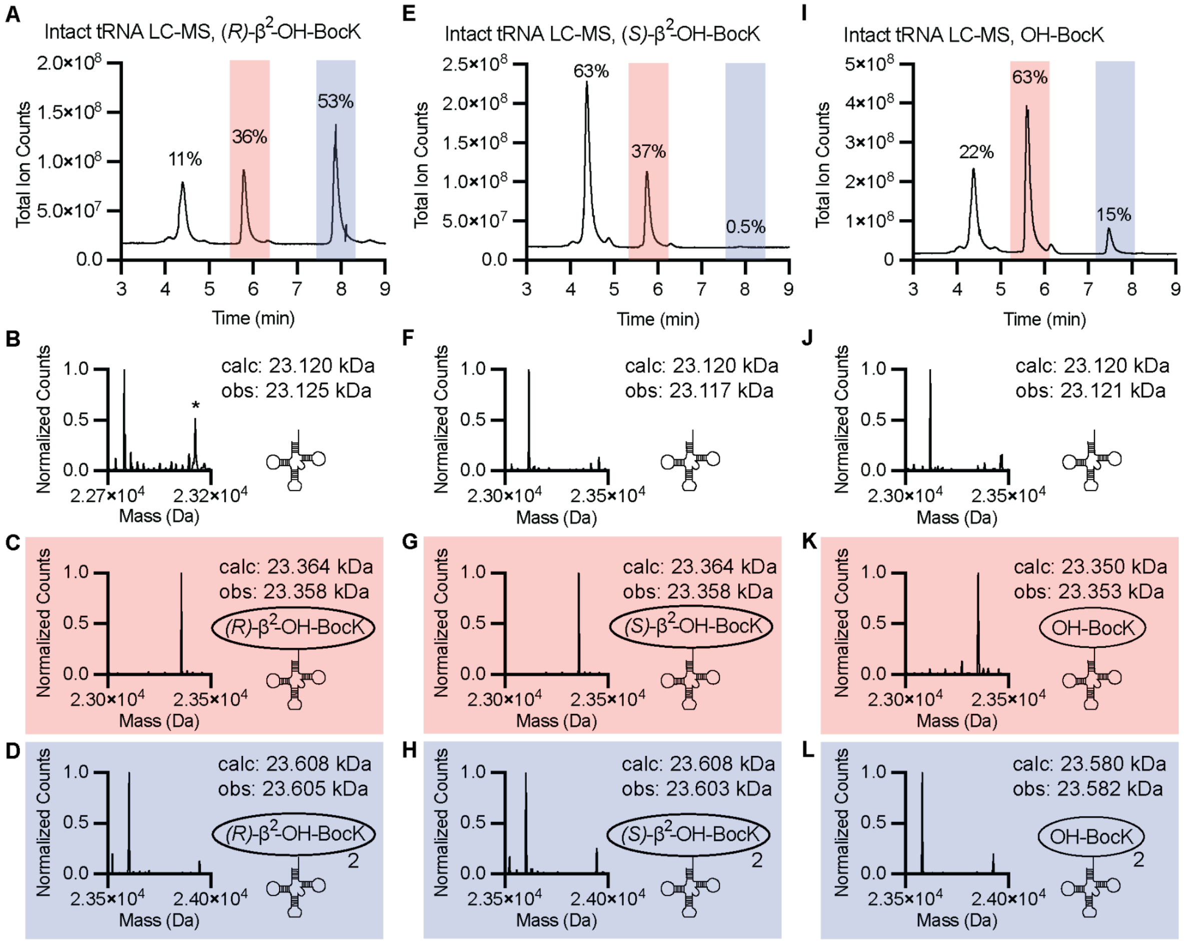

**Figure S9: Intact tRNA LC-MS of the products of *in vitro* tRNA$^{Pyl}$ acylations with various hydroxy-acid substrates confirms presence of unreacted tRNA$^{Pyl}$, monoacyl, and diacyl-tRNA$^{Pyl}$.** For all reactions, peaks in white correspond to unreacted tRNA$^{Pyl}$, peaks in red correspond to monoacyl-tRNA$^{Pyl}$, and peaks in blue correspond to diacyl-tRNA$^{Pyl}$. **(A)** Shown is the TIC of tRNA$^{Pyl}$ purified from an aminoacylation reaction containing 25 µM tRNA$^{Pyl}$, 12.5 µM *M. alvus* PylRS, and 10 mM (*R*)-β$^2$-OH-BocK incubated for 2 h at 37°C. Shown are deconvoluted mass spectra of **(B)** unreacted tRNA$^{Pyl}$ **(C)** (*R*)-β$^2$-OH-BocK-tRNA$^{Pyl}$ and **(D)** di-(*R*)-β$^2$-OH-BocK-tRNA$^{Pyl}$ derived from the highlighted peaks in **(A). (E)** Shown is the TIC of tRNA$^{Pyl}$ purified from an aminoacylation reaction containing 25 µM tRNA$^{Pyl}$, 12.5 µM *M. alvus* PylRS, and 10 mM (*S*)-β$^2$-OH-BocK incubated for 2 h at 37°C. Shown are deconvoluted mass spectra of **(F)** unreacted tRNA$^{Pyl}$ **(G)** (*S*)-β$^2$-OH-BocK-tRNA$^{Pyl}$ and **(H)** di-(*S*)-β$^2$-OH-BocK-tRNA$^{Pyl}$ derived from the highlighted peaks in **(E). (I)** Shown is the TIC of tRNA$^{Pyl}$ purified from an aminoacylation reaction containing 25 µM tRNA$^{Pyl}$, 2.5 µM *M. alvus* PylRS, and 10 mM OH-BocK incubated for 2 h at 37°C. Shown are deconvoluted mass spectra of **(J)** unreacted tRNA$^{Pyl}$ **(K)** OH-BocK-tRNA$^{Pyl}$ and **(L)** di-OH-BocK-tRNA$^{Pyl}$ derived from the highlighted peaks in **(I).**

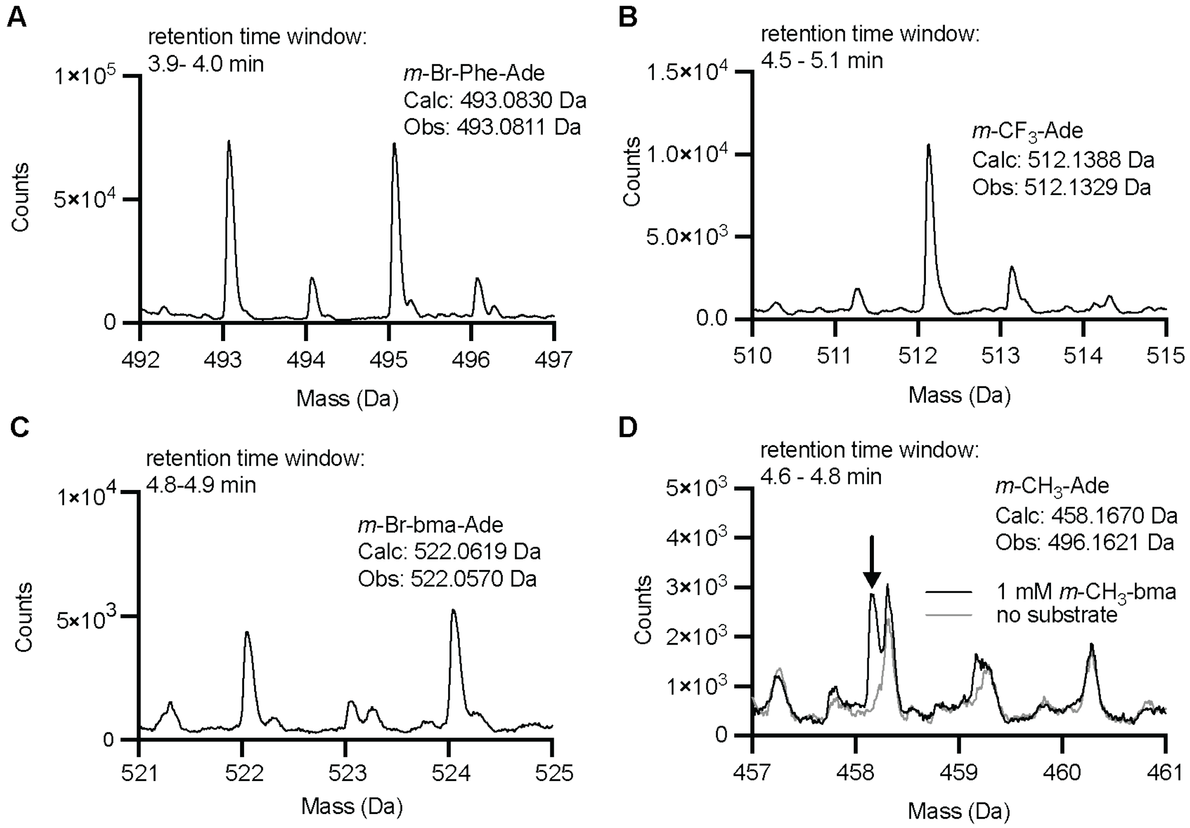

**Figure S10:** *In vivo* acylation with benzylmalonate derivatives is detectable with PARTI. All samples originated from C321 *E. coli* cells expressing *Ma*FRSA and *Ma*tRNA$^{Pyl}$ and grown with 1 mM respective substrate, and all calculated masses are [M+H]. Experimental details are identical to those in Figure 6. **(A)** The mass spectrum of *m*-Br-Phe-Ade detected over 3.9-4.0 min after PARTI with cells grown with 1 mM *m*-Br-Phe. **(B)** The mass spectrum of *m*-$CF_3$-bma-Ade detected over 4.5-5.1 min after PARTI with cells grown with 1 mM *m*-$CF_3$-bma. **(C)** The mass spectrum of *m*-Br-bma-Ade detected over 4.8-4.9 min after PARTI with cells grown with 1 mM *m*-Br-bma. **(D)** Overlaid mass spectra detected over 4.5-5.1 min after PARTI with cells grown without substrate (gray) or 1 mM *m*-$CH_3$-bma (black) highlighting the appearance of *m*-$CH_3$-bma-Ade only in the 1 mM *m*-$CH_3$-bma condition.

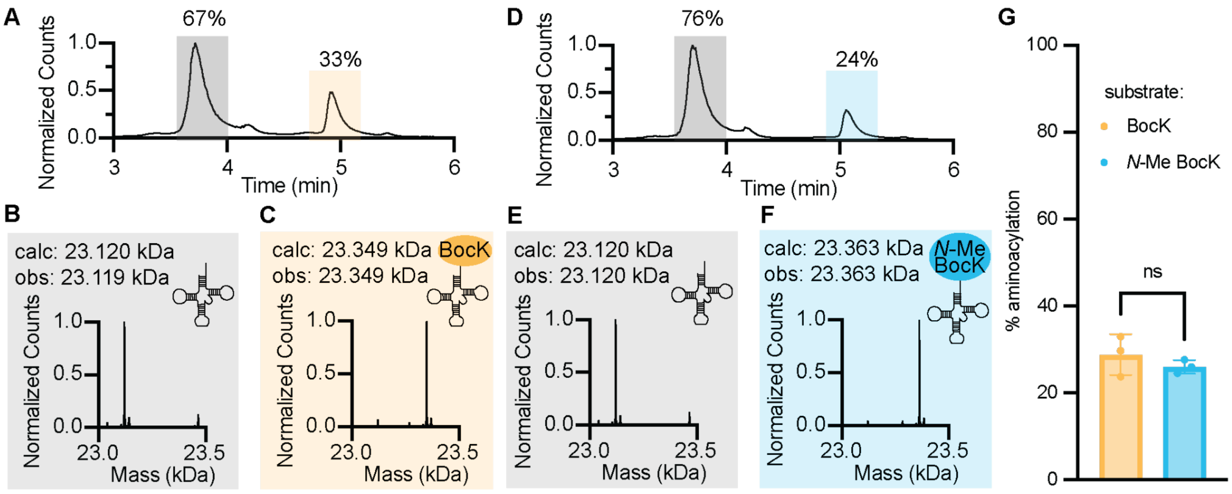

**Figure S11: BocK and *N*-Me BocK display similar *in vitro* activity with PylRS and tRNA$^{Pyl}$**
**(A)** Intact tRNA LC-MS TIC showing the products of an aminoacylation reaction containing 5 µM *M. alvus* PylRS, 25 µM tRNA$^{Pyl}$, and 10 mM BocK incubated for 2 h at 37°C. TIC signal is normalized to the highest signal in the trace. The peak highlighted in gray corresponds to unreacted tRNA$^{Pyl}$ and the peak in yellow corresponds to BocK-tRNA$^{Pyl}$. Deconvoluted mass spectra of **(B)** unreacted tRNA$^{Pyl}$ or **(C)** BocK-tRNA$^{Pyl}$ derived from the highlighted peaks in **(A).** The respective expected and observed masses of each product are shown. **(D)** Intact tRNA LC-MS TIC showing the products of an aminoacylation reaction containing 5 µM PylRS, 25 µM tRNA$^{Pyl}$, and 10 mM BocK incubated for 2 h at 37°C. TIC signal is normalized to the highest signal in the trace. The peak highlighted in gray corresponds to unreacted tRNA$^{Pyl}$ and the peak in blue corresponds to *N*-Me BocK-tRNA$^{Pyl}$. Deconvoluted mass spectra of **(E)** unreacted tRNA$^{Pyl}$ or **(C)** *N*-Me BocK-tRNA$^{Pyl}$ derived from the highlighted peaks in **(F).** The respective expected and observed masses of each product are shown. (**G**) Percent acylation determined by intact tRNA LC-MS for aminoacylation reactions containing 5 µM PylRS, 25 µM tRNA$^{Pyl}$, and either 10 mM BocK or 10 mM *N*-Me BocK incubated for 2 h at 37°C. Reactions were carried out in technical triplicate, with % acylation yields for BocK-tRNA$^{Pyl}$ (yellow, mean = 28.77, SD = 4.70) and *N*-Me-BocK-tRNA$^{Pyl}$ (blue, mean = 25.99, SD = 1.55) shown. Statistical analysis bars represent the results of a two-tailed unpaired t-test. $p > 0.05$ =ns; $p \leq 0.05$ = *; $p \leq 0.01$ = **; $p \leq 0.001$ = ***.

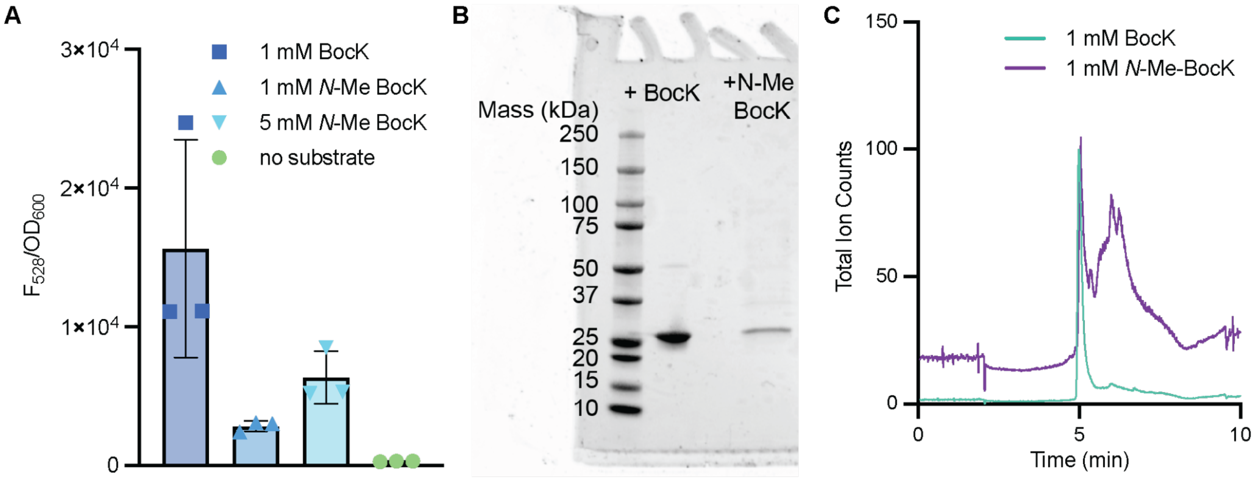

**Figure S12: Expression and purification of sfGFP-200TAG from cells supplemented with BocK or *N*-Me BocK. (A)** Bar graph comparing sfGFP fluorescence (ex: 485 nm em: 528 nm) normalized to cell optical density ($A_{600}$) after 24 hours of expression in the plate reader assay. *E. coli* Top10 cells expressing *Ma*PylRS, *Ma*tRNA$^{Pyl}$ and sfGFP-200TAG were supplemented with no substrate (lime, mean = 316.94 SD= 28.72 n = 3), 1 mM BocK (indigo, mean = 15660.94 SD = 7850.41 n= 3), 1 mM N-Me BocK (blue, mean = 2854.62 SD = 374.88 n=3), or 5 mM *N*-Me BocK (light blue, mean = 6360.60 SD = 1876.52 n=3). **(B)** SDS gel (Any kD™ Mini-PROTEAN® TGX™) of purified sfGFP-200TAG where *E. coli* Top10 cells were supplemented with 1 mM BocK (lane 2) or 1 mM *N*-Me BocK (lane 4). No sample is in lane 3 and lane 1 contains protein ladder (Precision Plus Protein™ Dual Color Standards, BioRad). **(C)** TICs following LC-HRMS analysis of protein samples shown in **(B).** The trace in teal is sfGFP-200TAG purified from cells grown with 1 mM BocK, and in purple is sfGFP-200TAG purified from cells grown with 1 mM *N*-Me BocK. The purple trace is shifted up 5 units for visibility. Because the sfGFP yield was lower in the 1 mM *N*-Me BocK condition, background signal from impurity is more intense relative to sfGFP, which eluted at 5 min.

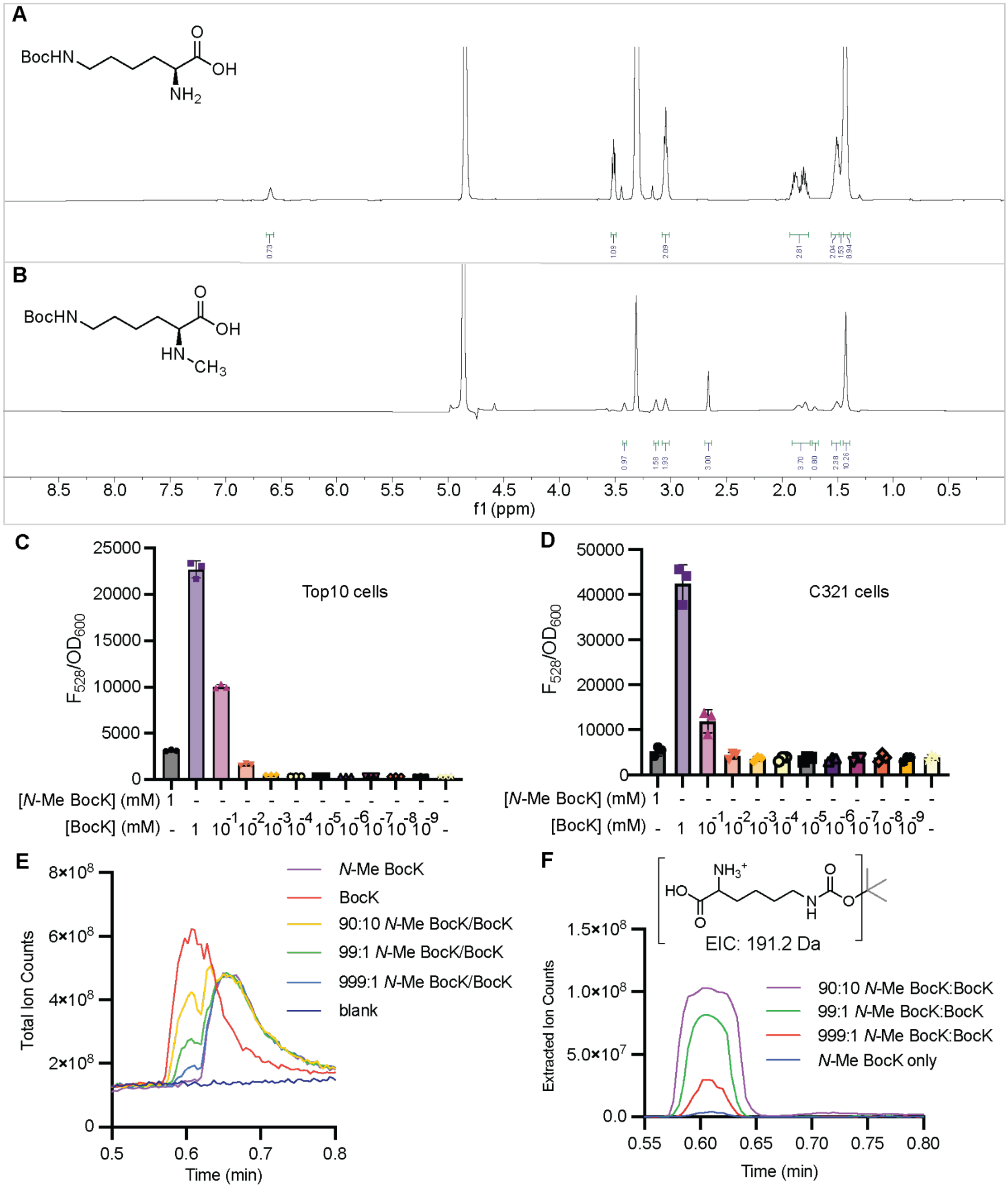

**Figure S13: Verification of high *N*-Me BocK purity and identification of BocK concentration required to reconstitute sfGFP signal observed from *E.* coli grown with 1 mM *N*-Me BocK.** 1H NMR spectra of (**A**) commercial BocK and (**B**) *N*-Me-BocK. Spectra were acquired on a Bruker 500 MHz NMR in methanol-*d4*. (**C**) sfGFP fluorescence at 528 nm over optical density ($OD_{600}$) of *E. coli* Top10 cells expressing PylRS, $tRNA^{Pyl}$, and sfGFP-200TAG with no substrate, 1mM *N*-Me BocK, or a serial dilution of BocK (1 mM - $1x10^{-9}$ mM)

measured 24 h after induction with 1 mM IPTG. (**D)** sfGFP fluorescence at 528 nm over optical density ($OD_{600}$) of *E. coli* C321 cells expressing PylRS, $tRNA^{Pyl}$, and sfGFP-200TAG with no substrate, 1mM *N*-Me BocK, or a serial dilution of BocK (1 mM - $1x10^{-9}$ mM) measured 24 h after induction with 1 mM IPTG. **(E)** TIC traces following small molecule MS analysis of solutions of *N*-Me BocK (purple), BocK (red), or *N*-Me BocK:BocK mixtures in ratios of 90:10 (yellow), 99:1 (green), and 999:1 (blue). A blank injection of only water is shown in indigo. **(F)** Overlaid EICs for the expected mass of a prominent BocK ion (calc [M+H]: 191.2 Da, fragment shown in brackets) following small molecule MS analysis of *N*-Me BocK (blue) and *N*-Me BocK:BocK mixtures in ratios of 90:10 (magenta), 99:1 (green), and 999:1 (red). Even 0.1% of a BocK impurity is detectable by mass spectrometry.

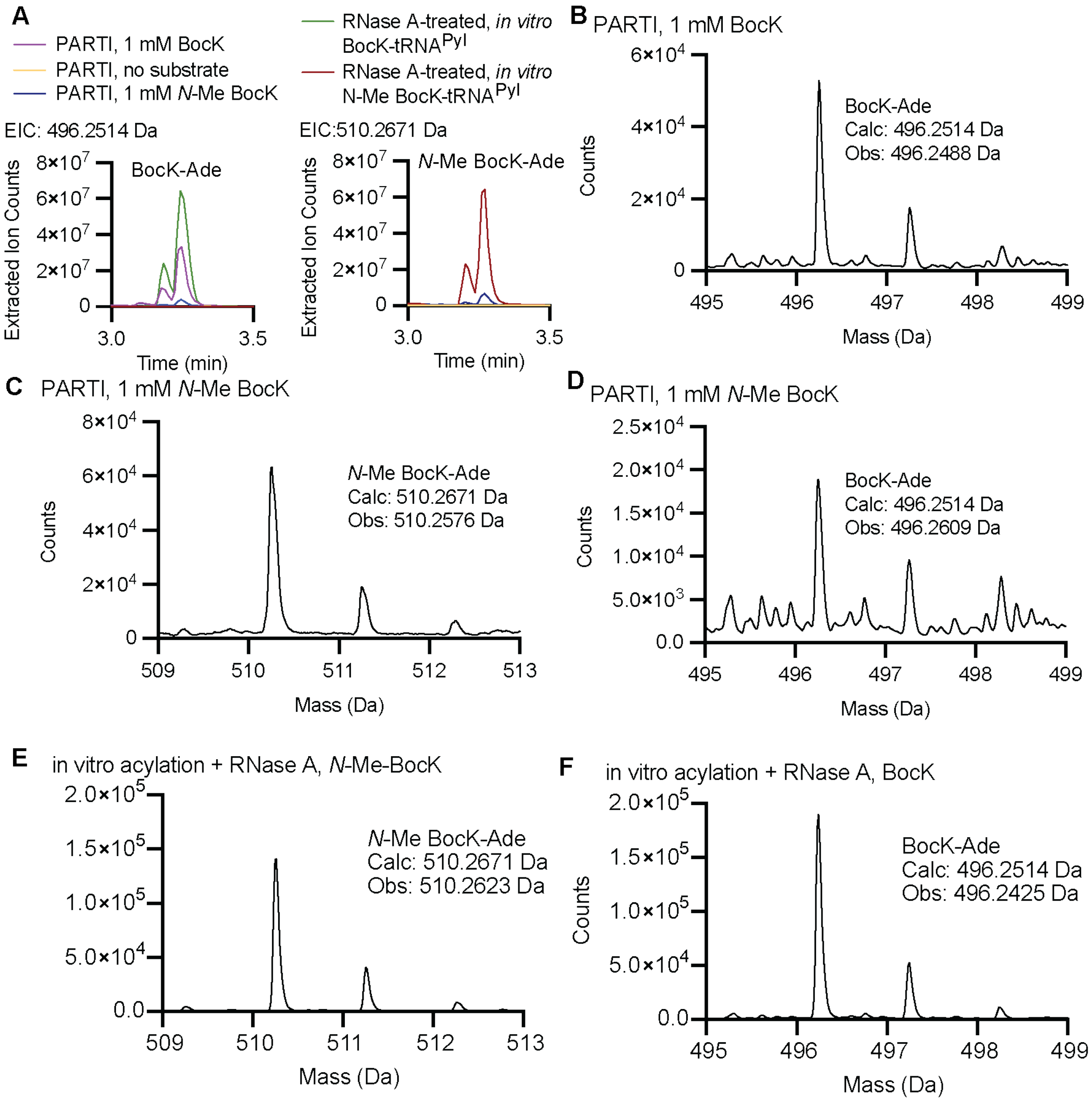

**Figure S14**: **Both BocK-Ade and *N*-Me BocK-Ade are detected following PARTI with cells supplemented only with *N*-Me BocK. (A)** Overlaid EICs of BocK-Ade, at left, and *N*-Me-BocK, at right, detected by LC-HRMS following either RNase A treatment of *in vitro* $tRNA^{Pyl}$ acylated with BocK (green) or *N*-Me-BocK (red) or following PARTI with *E. coli* DH5α cells expressing the *Ma*PylRS/$tRNA^{Pyl}$ pair and grown with 1 mM BocK (pink), no substrate (yellow) or 1 mM *N*-Me BocK (navy). LC-HRMS was carried out as described except for the length of run time and MS collection window. Chromatography was carried out with mobile phase B at 4% for 1.89 minutes followed by a linear gradient from 4 to 40% over 1.75 minutes. Then, mobile phase B underwent a gradient from 40 to 100% over 0.56 minutes and a subsequent gradient from 100 to 4% over 0.98 minutes. Mobile phase B was held at 4% for an additional

1.12 minutes. Mass spectrometry data were collected between 1.4 and 6.3 minutes. Acylations were performed by incubating 25 μM tRNA$^{Pyl}$, 5 μM PylRS, and 10 mM BocK or *N*-Me BocK for 3 h. **(B)** The mass spectrum of BocK-Ade detected following PARTI with *E. coli* Top10 cells expressing the *Ma*PylRS/tRNA$^{Pyl}$ pair and grown with 1 mM BocK. Shown are the mass spectra of **(C)** *N*-Me BocK-Ade and **(D)** BocK-Ade detected following PARTI with *E. coli* Top10 cells expressing the *Ma*PylRS/tRNA$^{Pyl}$ pair and grown with 1 mM *N*-Me BocK. **(E)** The mass spectrum of *N*-Me BocK-Ade detected following RNase A treatment of *in vitro* *N*-Me BocK-tRNA$^{Pyl}$. **(F)** The mass spectrum of BocK-Ade detected following RNase A treatment of *in vitro* BocK-tRNA$^{Pyl}$.

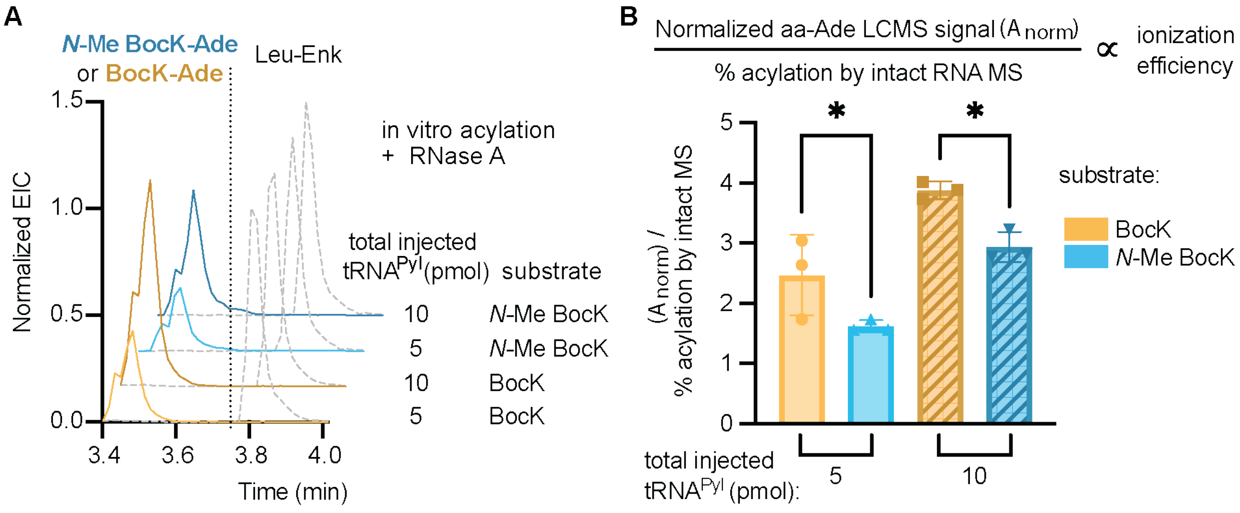

**Figure S15:** BocK and *N*-Me BocK RNase A cleavage products ionize differently by LC-HRMS, despite being acylated to $tRNA^{Pyl}$ at comparable levels *in vitro*. **(A)** Overlaid EICs from LC-HRMS analysis of *in vitro* $tRNA^{Pyl}$ acylated either with BocK or *N*-Me BocK and treated with RNase A. Injections containing 4 ng Leu-Enk and either 5 or 10 pmol of combined unreacted and acyl-$tRNA^{Pyl}$ were analyzed, and EICs for BocK-Ade (in dark and light gold, [M+H]: 496.2514 Da) or *N*-Me BocK-Ade (in dark and light blue, [M+H]: 510.2671 Da) are shown normalized to Leu-Enk (in gray dotted line, [M+H]: 556.2766 Da). **(B)** Comparison of relative ionization efficiencies between *N*-Me BocK-Ade and BocK-Ade. The equation shown describes that the peak area of the aa-Ade LC-HRMS signal ($A_{norm}$) following RNase A treatment of an *in vitro* $tRNA^{Pyl}$ acylation divided by the percent acylation determined by intact tRNA LC-MS is proportional to the ionization efficiency of the aa-Ade. Shown in the bar graph are aa-Ade LC-HRMS signals normalized to Leu-Enk ($A_{norm}$) divided by the percent acylation determined by intact tRNA LC-MS. *In vitro* acylations of either *N*-Me BocK or BocK onto $tRNA^{Pyl}$ were completed in technical triplicate, then, aliquots from each reaction containing 5 or 10 pmol total $tRNA^{Pyl}$ were treated with RNase A and analyzed by LC-HRMS with the Leu-Enk internal standard. $A_{norm}$ was determined as described. In yellow are samples where BocK was the substrate (solid yellow bar: 5 pmol RNase A-treated $tRNA^{Pyl}$, mean = 2.47 SD = 0.67 n = 3; yellow striped bar: 10 pmol RNase A-treated $tRNA^{Pyl}$, mean = 3.88 SD = 0.14 n = 3 ), and in blue are samples where N-Me BocK was the substrate (solid blue bar: 5 pmol RNase A-treated $tRNA^{Pyl}$, mean = 1.62 SD = 0.01 n = 3; blue striped bar: 10 pmol RNase A-treated $tRNA^{Pyl}$, mean = 2.94 SD = 0.25 n = 3 ). Acylations were carried out with 25 mM BocK or N-Me BocK, 5 µM PylRS, and 25 µm $tRNA^{Pyl}$ for 2 hours. LC-HRMS was carried out as described except for the length of run time and MS collection window. Chromatography was carried out with mobile phase B at 4% for 1.89 minutes followed by a linear gradient from 4 to 40% over 1.75 minutes. Then, mobile phase B underwent a gradient from 40 to 100% over 0.56 minutes and a subsequent gradient from 100 to 4% over 0.98 minutes. Mobile phase B was held at 4% for an additional 1.12 minutes. Mass spectrometry data were collected between 1.4 and 6.3 minutes.

**Table S1:** Oligomers and sequences used

| name | sequence |
|---|---|
| o-Pyl | BTN-<br>CGAACCCCGCTGGCTAGGTTTTAGAG |
| o-Phe | BTN-<br>CCGGACTCGGAATCGAACCAAGGACA<br>CGGGG |
| PylT transcription sequences (*in vitro*) | PylT-Fwd:<br>CTAATACGACTCACTATAGGGGGACGG<br>TCCGGCGACCAGCGGGTCTCTAAAACC<br>TAGCCA<br>PylT-Rev:<br>TGGCGAGAGACCGGGGCGTCGAACCC<br>CGCTGGCTAGGTTTTAGAGACCCGCTG<br>GTCGCCG |
| PheT transcription sequences (*in vitro*) | PheT-Fwd:<br>AATTCCTGCAGTAATACGACTCACTAT<br>AGCCCGGATAGCTCAGTCGGTAGAGC<br>AGG<br>PheT-Rev:<br>TGGTGCCCGGACTCGGAATCGAACCAA<br>GGACACGGGGATTTTCAATCCCCTGCT<br>CTA |
| PylT sequence – in pMega plasmid | GGGGGACGCGCAGCCTGGTAGCGCAG<br>CGCTAAAACCTAGCCAGCGGGGTTCGA<br>CGCCCCGGTCTCTCGCCA |
| PheT sequence (1) | GCCCGGATAGCTCAGTCGGTAGAGCA<br>GGGGATTGAAAATCCCCGTGTCCTTGG<br>TTCGATTCCGAGTCCGGGCACCA |

**Table S2**: Statistical information for all comparisons. All statistical tests were performed using GraphPad Prism 9 software. p > 0.05 = ns; p ≤ 0.05 = *; p ≤ 0.01 = **; p ≤ 0.001 = ***.

| Figure | Comparison | | **Mean Difference** | **95.00% CI** | **P-value** |
|---|---|---|---|---|---|
| Figure 3 | | | | | |
| | Ordinary one-way ANOVA, Sidak's multiple comparisons test, with a single pooled variance | | | | |
| | raw Phe-Ade day 1 vs. | raw Phe-Ade day 2 | 1268167 | 649522 to 1886813 | 0.0018 |
| | normalized Phe-Ade day 1 vs. | normalized Phe-Ade day 2 | -0.01906 | -618646 to 618646 | >0.9999 |
| | Ordinary one-way ANOVA, Dunnett's multiple comparisons test, with a single pooled variance | | | | |
| | 1x total RNA | no RNase A | 0.1882 | 0.07423 to 0.3022 | 0.001 |
| | | o-Pyl | 0.1882 | 0.07423 to 0.3022 | 0.001 |
| | | 2x bead | 0.1219 | 0.007883 to 0.2359 | 0.0332 |
| | | 2x oligo (o-Phe) | -0.1015 | -0.2155 to 0.01251 | 0.0939 |
| | | 0.5x RNA | 0.04263 | -0.07137 to 0.1566 | 0.8302 |

| | | | | | |
|---|---|---|---|---|---|
| | | 2x RNA | -0.1444 | -0.2584 to - 0.03043 | 0.01 |
| Figure 5 | Ordinary one-way ANOVA, Tukey's multiple comparisons test, with a single pooled variance | | | | |
| | OH-BocK Ade vs. | *(S)*-β$^{2}$-OH-BocK-Ade | 2.566 | 1.827 to 3.304 | <0.0001 |
| | | no substrate | 2.763 | 2.024 to 3.501 | <0.0001 |
| | | *(R)*-β$^{2}$-OH-BocK-Ade | 0.5856 | -0.1530 to 1.324 | 0.1548 |
| | *(S)*-β$^{2}$-OH-BocK-Ade vs. | *(R)*-β$^{2}$-OH-BocK-Ade | -1.98 | -2.719 to - 1.241 | <0.0001 |
| | | no substrate | 0.1969 | -0.5417 to 0.9355 | 0.9404 |
| | *(R)*-β$^{2}$-OH-BocK-Ade vs. | no substrate | 2.177 | 1.438 to 2.916 | <0.0001 |
| | diacyl *(R)*-β$^{2}$-OH-BocK-Ade vs. | no substrate | 0.1353 | -0.6033 to 0.8739 | 0.9877 |
| Supplementary Figure 8 | | | | | |
| | Ordinary one-way ANOVA, Tukey's multiple comparisons test, with a single pooled variance | | | | |
| | no substrate aa-Ade vs. | OH-BocK-Ade | -1.231 | -1.503 to - 0.9585 | 0.0002 |
| | | *(R)*-β$^{2}$-OH-BocK-Ade | -1.076 | -1.348 to - 0.8041 | 0.0003 |

| | | | | | |
|---|---|---|---|---|---|
| | | (*S)*-β$^2$-OH-BocK-Ade | -0.06063 | -0.3328 to 0.2115 | 0.8035 |
| | OH-BocK-Ade vs. | (*R)*-β$^2$-OH-BocK-Ade | 0.1545 | -0.1177 to 0.4266 | 0.2386 |
| | | (*S)*-β$^2$-OH-BocK-Ade | 1.17 | 0.8979 to 1.442 | 0.0002 |
| | (*R)*-β$^2$-OH-BocK-Ade vs. | (*S)*-β$^2$-OH-BocK-Ade | 1.016 | 0.7434 to 1.288 | 0.0004 |
| | Ordinary one-way ANOVA, Tukey's multiple comparisons test, with a single pooled variance | | | | |
| | Ade w no substrate vs. | Ade w added OH-BocK | 1.158 | 0.5542 to 1.762 | 0.005 |
| | | Ade w added (*R)*-β$^2$-OH-BocK | 1.115 | 0.5105 to 1.719 | 0.0058 |
| | | Ade w added (*S)*-β$^2$-OH-BocK | -0.0653 | -0.6694 to 0.5388 | 0.9682 |
| | Ade w added OH-BocK vs. | Ade w added (*R)*-β$^2$-OH-BocK | -0.04369 | -0.6478 to 0.5604 | 0.9898 |
| | | Ade w added (*S)*-β$^2$-OH-BocK | -1.224 | -1.828 to -0.6195 | 0.0041 |
| | Ade w added (*R)*-β$^2$-OH-BocK vs. | Ade w added (*S)*-β$^2$-OH-BocK | -1.18 | -1.784 to -0.5758 | 0.0047 |
| Figure 6 | Ordinary one-way ANOVA, Tukey's multiple comparisons test, with a single pooled variance | | | | |
| | No substrate vs. | *m*-$CF_3$-bma-Ade | -0.2335 | -0.3471 to -0.1199 | 0.0005 |

| | | | | | |
|---|---|---|---|---|---|
| | | *m*-Br-bma-Ade | -0.04106 | -0.1427 to 0.06053 | 0.6653 |
| | | *m*-$CH_3$-bma-Ade | -0.02546 | -0.1270 to 0.07613 | 0.9104 |
| | | *m*-Br-Phe-Ade | -0.6424 | -0.7440 to -0.5409 | <0.0001 |
| | *m*-$CF_3$-bma-Ade vs. | *m*-Br-bma-Ade | 0.1924 | 0.07883 to 0.3060 | 0.0021 |
| | | *m*-$CH_3$-bma-Ade | 0.208 | 0.09444 to 0.3216 | 0.0012 |
| | | *m*-Br-Phe-Ade | -0.409 | -0.5226 to -0.2954 | <0.0001 |
| | *m*-Br-bma-Ade vs. | *m*-$CH_3$-bma-Ade | 0.0156 | -0.08599 to 0.1172 | 0.9835 |
| | | *m*-Br-Phe-Ade | -0.6014 | -0.7030 to -0.4998 | <0.0001 |
| | *m*-CH3-bma-Ade vs | *m*-Br-Phe-Ade | -0.617 | -0.7186 to -0.5154 | <0.0001 |
| Figure 7 | Ordinary one-way ANOVA, Tukey's multiple comparisons test, with a single pooled variance | | | | |
| | BocK-Ade, 1 mM BocK vs. | *N*-Me BocK-Ade, 1 mM BocK | 1.788 | 0.8001 to 2.776 | 0.0004 |

| | | | | | |
|---|---|---|---|---|---|
| | | BocK-Ade, 1 mM N-Me BocK | 1.541 | 0.5525 to 2.529 | 0.0016 |
| | | *N*-Me BocK-Ade, 1 mM *N*-Me BocK | 1.222 | 0.2338 to 2.210 | 0.0109 |
| | | BocK-Ade, no substrate | 1.788 | 0.8001 to 2.776 | 0.0004 |
| | | *N*-Me BocK-Ade, no substrate | 1.788 | 0.8001 to 2.776 | 0.0004 |
| | N-Me BocK-Ade, 1 mM BocK vs. | BocK-Ade, 1 mM *N*-Me BocK | -0.2476 | -1.236 to 0.7405 | 0.9992 |
| | | *N*-Me BocK-Ade, 1mM *N*-Me BocK | -0.5663 | -1.554 to 0.4219 | 0.5999 |
| | | BocK-Ade, no substrate | 0 | -0.9881 to 0.9881 | >0.9999 |
| | | *N*-Me BocK-Ade, no substrate | 0 | -0.9881 to 0.9881 | >0.9999 |
| | BocK-Ade, 1 mM N-Me BocK vs. | *N*-Me BocK-Ade, 1 mM *N*-Me BocK | -0.3187 | -1.307 to 0.6695 | 0.9899 |
| | | BocK-Ade, no substrate | 0.2476 | -0.7405 to 1.236 | 0.9992 |
| | | *N*-Me BocK-Ade, no substrate | 0.2476 | -0.7405 to 1.236 | 0.9992 |
| | *N*-Me BocK-Ade, 1 mM N-Me BocK vs. | BocK-Ade, no substrate | 0.5663 | -0.4219 to 1.554 | 0.5999 |
| | | *N*-Me BocK-Ade, no substrate | 0.5663 | -0.4219 to 1.554 | 0.5999 |

| | | | | | |
|---|---|---|---|---|---|
| | BocK-Ade w no substrate vs. | *N*-Me BocK-Ade, no substrate | 0 | -0.9881 to 0.9881 | >0.9999 |
| Supplementary Figure 11 | | | | | |
| S11G | Unpaired two-tailed t test | | | | |
| | % aminoacylation, BocK vs. *N*-Me BocK | | -2.773 ± 2.859 | -10.71 to 5.164 | 0.387 |
| S11H | Ordinary one-way ANOVA, Tukey's multiple comparisons test, with a single pooled variance | | | | |
| | no substrate vs. | 1 mM BocK | -15344 | -25908 to -4780 | 0.0071 |
| | | 1 mM *N*-Me BocK | -2538 | -13102 to 8026 | 0.8661 |
| | | 5 mM *N*-Me BocK | -6044 | -16608 to 4520 | 0.3266 |
| | 1 mM BocK vs. | 1 mM *N*-Me BocK | 12806 | 2242 to 23370 | 0.0195 |
| | | 5 mM *N*-Me BocK | 9300 | -1264 to 19864 | 0.0859 |
| | 1 mM *N*-Me BocK vs. | 5 mM *N*-Me BocK | -3506 | -14070 to 7058 | 0.7199 |
| Supplementary Figure 15 | Ordinary one-way ANOVA, Sidak's multiple comparisons test, with a single pooled variance | | | | |
| | aa-Ade/% acylation, 5 pmol BocK acylation vs. | 5 pmol *N*-Me BocK acylation | 0.8531 | 0.02936 to 1.677 | 0.0431 |

| | | | | | |
|---|---|---|---|---|---|
| | aa-Ade/% acylation, 10 pmol BocK acylation vs. | 10 pmol *N*-Me BocK acylation | 0.9417 | 0.1180 to 1.765 | 0.0276 |

**Table S3:** Major ions and peak areas for *in vitro* tRNA acylations analyzed using intact tRNA LC-MS.

| | major ion (m/z) | area | % of total |
|---|---|---|---|
| **Figure 2** | | | |
| unreacted $tRNA^{Phe}$ | 881.4094 | 533715 | 39.07 |
| Phe-$tRNA^{Phe}$ | 886.6588 | 832253 | 60.93 |
| **Figure 4** | | | |
| unreacted $tRNA^{Pyl}$ | 824.6992 | 416824 | 63.03 |
| BocK-$tRNA^{Pyl}$ | 832.8477 | 244472 | 36.97 |
| **Supplementary Figure 5** | | | |
| unreacted $tRNA^{Pyl}$ | 824.6339 | 316143 | 35.30 |
| OH-BocK-$tRNA^{Pyl}$ | 832.8148 | 457569 | 51.10 |
| di-OH-BocK-$tRNA^{Pyl}$ | 841.0295 | 121752 | 13.60 |
| **Supplementary Figure 9** | | | |
| unreacted $tRNA^{Pyl}$ | 824.7122 | 1523085 | 22.20 |
| OH-BocK-$tRNA^{Pyl}$ | 832.9001 | 4345872 | 63.34 |
| di-OH-BocK-$tRNA^{Pyl}$ | 841.1154 | 992169 | 14.46 |
| unreacted tRNAPyl | 855.0233 | 261180 | 11.08 |
| (*R)*-$\beta^2$-OH-BocK-$tRNA^{Pyl}$ | 864.0286 | 846021 | 35.90 |
| di-(*R)*-$\beta^2$-OH-BocK-$tRNA^{Pyl}$ | 873.0341 | 1249130 | 53.01 |

| | | | |
|---|---|---|---|
| unreacted tRNA$^{Pyl}$ | 855.0233 | 1949446 | 62.76 |
| *(S)*-β$^2$-OH-BocK-tRNA$^{Pyl}$ | 864.0286 | 1141183 | 36.74 |
| di-*(S)*-β$^2$-OH-BocK-tRNA$^{Pyl}$ | 873.0341 | 15463 | 0.48 |
| **Supplementary Figure 11** | | | |
| unreacted tRNA$^{Pyl}$ | 855.2896 | 3525875 | 67.03 |
| BocK-tRNA$^{Pyl}$ | 832.8555 | 1733719 | 32.96 |
| unreacted tRNA$^{Pyl}$ | 855.2829 | 3577458 | 70.35 |
| BocK-tRNA$^{Pyl}$ | 832.8489 | 1507876 | 29.65 |
| unreacted tRNA$^{Pyl}$ | 855.2896 | 4445625 | 76.32 |
| BocK-tRNA$^{Pyl}$ | 832.8555 | 1379681 | 23.68 |
| unreacted tRNA$^{Pyl}$ | 855.2896 | 4715029 | 75.52 |
| *N*-Me-BocK-tRNA$^{Pyl}$ | 833.3456 | 1528511 | 24.48 |
| unreacted tRNA$^{Pyl}$ | 855.2896 | 3883568 | 72.42 |
| *N*-Me-BocK-tRNA$^{Pyl}$ | 833.3546 | 1479076 | 27.58 |
| unreacted tRNA$^{Pyl}$ | 855.2829 | 4094939 | 74.08 |
| *N*-Me-BocK-tRNA$^{Pyl}$ | 833.348 | 1432538 | 25.92 |
